# Integrative Multi-cohort Transcriptomics and Network Pharmacology Analysis Reveals Key Network Nodes and Potential Drug Clues in PCOS Granulosa Cells

**DOI:** 10.64898/2026.04.01.715808

**Authors:** Xuan Zhang, Jie Fang, Zhinan Liu, Shuangsang Li, Fanhui Jin, Linlin Guo, Ruonan Qiang, Yinan Zhu, Tingting Hou, Jingjing Li, Yanfeng Liu

## Abstract

**Background:** Polycystic ovary syndrome (PCOS) is a prevalent endocrine disorder with complex pathophysiology and limited therapeutic options. Identifying key molecular drivers and potential drug candidates is critical for improving clinical outcomes.

**Methods:** We integrated multi-cohort transcriptomics (GSE155489, GSE138518, GSE226146) with weighted gene co-expression network analysis (WGCNA), protein-protein interaction (PPI) network analysis, and drug repurposing. Differential expression analysis identified 1,039 DEGs, and WGCNA identified 10 PCOS-associated modules. Intersection of DEGs with module genes yielded 498 core candidate genes, which were subjected to functional enrichment, PPI network analysis, and connectivity map-based drug repurposing (CLUE/LINCS). Candidate drugs were further evaluated by molecular docking and ADMET prediction using a triple intersection strategy (hub genes, high differential expression, drug-target evidence).

**Results:** Functional enrichment revealed significant enrichment in cell adhesion and TGF-beta signaling. PPI network analysis identified CD44 as the top hub gene (degree=42). Drug repurposing identified 106 candidate drugs, including troglitazone and enzalutamide. Using the triple intersection strategy, five genes (ID2, NR4A1, GJA5, ID1, MYH11) were prioritized for molecular docking. GJA5 showed strong predicted binding affinity with flufenamic acid (−7.88 kcal/mol), and cytosporone B exhibited favorable druglikeness (0 Lipinski violations).

**Conclusion:** This study systematically characterizes PCOS-associated gene networks and provides a prioritized set of candidate targets and drugs through a purely computational framework. CD44 emerges as a key network node with potential relevance in PCOS pathophysiology. These findings offer testable hypotheses for future mechanistic studies and drug discovery efforts in PCOS.

## 1. Introduction

Polycystic ovary syndrome (PCOS) is the most common endocrine and metabolic disorder in women of reproductive age, with a global prevalence of approximately 5-10%[1]. The clinical presentation of PCOS is highly heterogeneous, with main features including hyperandrogenism, ovulatory dysfunction, and polycystic ovarian morphology, often accompanied by insulin resistance, obesity, metabolic syndrome, and increased cardiovascular disease risk[2,33]. PCOS is one of the leading causes of female infertility, accounting for approximately 70% of anovulatory infertility[3]. Furthermore, even when PCOS patients achieve pregnancy, they face elevated risks of pregnancy complications, including gestational diabetes, preeclampsia, and preterm birth[4]. Therefore, understanding the molecular mechanisms of PCOS and identifying key pathogenic gen es and signaling pathways are of great importance for improving reproductive outcomes and long-term health in PCOS patients.

The pathophysiology of PCOS is complex, involving functional abnormalities across multiple organ systems. At the ovarian level, granulosa cells in PCOS patients exhibit aberrant proliferation, differentiation, and steroid hormone synthesis[5]. Granulosa cells are critical cell types in follicular development, responsible for estrogen synthesis, oocyte nutritional support, and ovulation signaling. Abnormal expression of multiple genes in PCOS granulosa cells, including those involved in cell cycle regulation, inflammatory response, and metabolic pathways, leads to follicular developmental arrest and ovulatory dysfunction[6]. At the metabolic level, approximately 50-70% of PCOS patients exhibit insulin resistance, which not only affects glucose and lipid metabolism but also exacerbates hyperandrogenism by enhancing androgen synthesis in the ovaries and adrenal glands[7]. At the endometrial level, PCOS patients demonstrate reduced endometrial receptivity, characterized by abnormal endometrial proliferation, impaired decidualization, and enhanced inflammatory response, which are important contributors to PCOS-associated infertility and pregnancy loss[8].

Despite significant progress in PCOS research, its molecular mechanisms remain incompletely elucidated. Traditional candidate gene approaches often focus on single or a few genes, making it difficult to comprehensively reveal the complex pathological network of PCOS. In recent years, the development of high-throughput transcriptomic sequencing technologies has provided powerful tools for systematic investigation of PCOS molecular mechanisms[9,34]. By integrating transcriptomic data from multiple independent cohorts, genes with consistent alterations across different tissues (such as granulosa cells and endometrium) can be identified, improving the reliability and reproducibility of research findings[10,35]. Weighted gene co-expression network analysis (WGCNA) is a systems biology approach that constructs gene co-expression networks to identify gene modules highly correlated with disease phenotypes, thereby discovering key pathogenic genes and signaling pathways[11]. Protein-protein interaction (PPI) network analysis can further identify Hub genes within the network, which often play central regulatory roles in disease development and progression[12].

Drug repurposing is an efficient drug development strategy that screens approved or clinical-stage drugs to identify their potential applications in new indications[13]. Compared to traditional de novo drug development, drug repurposing offers advantages including lower cost, shorter timeline, and known safety profiles. The Connectivity Map (CMap) and Library of Integrated Network-Based Cellular Signatures (LINCS) databases catalog the effects of thousands of small molecule compounds on cellular gene expression, enabling identification of candidate drugs capable of reversing disease phenotypes by comparing disease gene expression signatures with drug-induced gene expression changes[14]. Molecular docking technology can predict the binding modes and affinities between small molecule drugs and target proteins, providing structural biology insights into drug-target interactions[15]. Combined with ADMET (Absorption, Distribution, Metabolism, Excretion, Toxicity) property predictions, the druglikeness of candidate compounds can be assessed, providing reference for subsequent experimental studies and further validation[16].

Based on the above background, this study aims to identify key genes and signaling pathways associated with PCOS and screen candidate targets and potential intervention clues by integrating multi-cohort transcriptomics and network pharmacology approaches. Specific objectives include: (1) identifying core genes associated with PCOS using differential expression analysis and WGCNA based on three independent cohorts from the GEO database (GSE155489, GSE138518, GSE226146); (2) identifying Hub genes through PPI network analysis to reveal key regulatory nodes in PCOS; (3) elucidating core pathological pathways of PCOS using functional enrichment analysis; (4) performing drug repurposing analysis based on the CLUE platform (Connectivity Map/LINCS database) to screen candidate therapeutic drugs; (5) evaluating drug-target interactions and druglikeness through molecular docking and ADMET predictions. This study can provide a traceable candidate list and analytical reference for PCOS-related mechanistic research and subsequent validation.

## 2. Materials and Methods

### 2.1 Data Sources and Preprocessing

This study utilized three publicly available datasets from the Gene Expression Omnibus (GEO) database (https://www.ncbi.nlm.nih.gov/geo/). GSE155489 consisted of granulosa cell RNA-seq count matrices (8 samples: 4 PCOS vs 4 controls), with sample grouping determined after consistency verification by reproducing published differential results from the original study: PCOS group included GC_B7, GC_B8, GC_B15, GC_B16; control group included GC_B13, GC_B14, GC_B2, GC_B30. GSE138518 consisted of granulosa cell RNA-seq expression matrices (6 samples: 3 PCOS: P14, P15, P16; 3 controls: N20, N21, N25). GSE226146 consisted of endometrial RNA-seq expression matrices (6 samples: 3 PCOS: PCOS1, PCOS2, PCOS3; 3 controls: OVULATE1, OVULATE2, OVULATE3).

The data preprocessing workflow was as follows: (1) downloading expression matrices and sample information from the GEO database; (2) performing normalization and variance-stabilizing transformation on the GSE155489 count matrix using DESeq2[27] for differential analysis and visualization; (3) organizing and filtering gene annotations for the FPKM matrices of GSE138518 and GSE226146, followed by log2(FPKM+1) transformation for differential analysis and downstream modeling; (4) gene filtering: retaining genes expressed in at least 50% of samples (expression value >0). Quality control employed principal component analysis (PCA) and hierarchical clustering analysis to assess the rationality of sample grouping and overall data quality.

### 2.2 Differential Expression Analysis

Differential expression analysis employed two strategies: DESeq2 was used for count data (GSE155489)[27]; limma was used for log2-transformed FPKM data (GSE138518 and GSE226146)[17]. The screening criteria for differentially expressed genes were: |log2 Fold Change (log2FC)| > 1 and adjusted P-value (adj.P.Val) < 0.05, with multiple testing correction performed using the Benjamini-Hochberg method. Hierarchical clustering analysis was performed on differentially expressed genes, heatmaps were generated to display gene expression patterns, and volcano plots were used to visualize the distribution of differentially expressed genes.

### 2.3 Weighted Gene Co-expression Network Analysis

Weighted gene co-expression network analysis (WGCNA) was performed on preprocessed genes from the GSE155489 dataset using the WGCNA package (version 1.72-1)[10]. The analysis workflow was as follows: (1) sample clustering to detect outlier samples; (2) selection of soft threshold power through the scale-free topology fit index, requiring R²>0.85; (3) identification of gene modules using the dynamic tree cut method, with minimum module size set to 30 and correlation coefficient threshold for merging similar modules set to 0.25; (4) calculation of module eigengenes (ME), representing the first principal component of gene expression within modules; (5) calculation of module-PCOS phenotype correlations using Pearson correlation coefficients, with P≤0.1 as the reference threshold for screening; (6) calculation of gene significance (GS) and module membership (MM), screening for highly significant module genes (P<0.01).

### 2.4 Core Candidate Gene Selection

An intersection strategy was employed to integrate results from differential expression analysis and WGCNA analysis, screening for core candidate genes that simultaneously satisfied differential expression significance and co-expression network relevance. Specifically, the intersection of GSE155489 differentially expressed genes (1,039 DEGs) and WGCNA highly significant module genes (3,777 module genes from MEred, MEgreen, and MEorangered3 modules, P<0.01) yielded 498 core candidate genes. This strategy ensured that candidate genes simultaneously reflected differential expression signals and co-expression network correlations, enhancing the interpretability and robustness of the candidate set to some extent.

### 2.5 Functional Enrichment Analysis

Functional enrichment analysis was performed on core candidate genes using the clusterProfiler package (version 4.6.0)[18]. The analysis included: (1) Gene Ontology (GO) enrichment analysis, covering three categories: Biological Process (BP), Cellular Component (CC), and Molecular Function (MF); (2) Kyoto Encyclopedia of Genes and Genomes (KEGG) pathway enrichment analysis; (3) Reactome pathway enrichment analysis. Enrichment analysis employed hypergeometric testing, with a significance threshold of adjusted P-value (adj.P) < 0.05, and multiple testing correction was performed using the Benjamini-Hochberg method. Bubble plots and bar charts were generated using the ggplot2 package to visualize enrichment results.

### 2.6 Protein-Protein Interaction Network Analysis

Based on the STRING database (https://string-db.org/, version 11.5), a protein-protein interaction (PPI) network was constructed for core candidate genes, with the interaction confidence threshold set to 0.4 (medium confidence). The PPI network (Figure 5A) contained 271 nodes and 974 edges, with a network average degree of 7.19 and network density of 0.027, indicating extensive interaction relationships among candidate genes. Network topology parameters were calculated using the NetworkAnalyzer plugin of Cytoscape software (version 3.9.0)[28], including degree, betweenness centrality, closeness centrality, and eigenvector centrality.

### 2.7 Drug Repurposing Analysis

Drug repurposing analysis was performed based on the CLUE platform (Connectivity Map/LINCS database) to identify candidate drugs capable of reversing PCOS gene expression patterns. The upregulated genes (272 genes) and downregulated genes (226 genes) from the 498 intersected genes were submitted to the CLUE platform as the PCOS disease expression signature for query. Drug screening criteria were: (1) connectivity score < -0.5, indicating the drug’s ability to reverse PCOS gene expression patterns; (2) tested in at least 2 cell lines; (3) priority given to drugs targeting Hub genes.

### 2.8 Molecular Docking Analysis

#### 2.8.1 Target Gene Selection Strategy

To select targets with greater operational feasibility for molecular docking evaluation, we employed a multi-dimensional integration strategy. This strategy comprehensively considered three key dimensions: network topological importance of genes, disease relevance, and drug accessibility.

Selection Criteria: Candidate genes must simultaneously satisfy the following three conditions:

1. Network Importance: Genes possess Hub gene characteristics in the protein-protein interaction (PPI) network. Hub genes refer to genes with high centrality metrics (degree, betweenness, closeness, eigenvector centrality) in the network, occupying key positions in the disease network, and their functional abnormalities may affect multiple downstream pathways.
2. Disease Relevance: Genes exhibit high differential expression in PCOS patients (|log2 fold change| > 2.0). The significance of differential expression reflects the magnitude of gene changes in disease states, with |logFC| > 2.0 indicating expression changes exceeding 4-fold, possessing strong biological directionality. Genes with high differential expression may be closer to key disease-related changes and are therefore more suitable for inclusion in subsequent target prioritization assessment.
3. Drug Accessibility: Genes are targeted by at least one candidate drug. This condition enables subsequent structure-based evaluation (such as molecular docking) to have operational candidate targets.

Scientific Rationale: This multi-dimensional integration strategy is based on the following considerations: (1) Hub genes play key roles in disease networks, but not all Hub genes are directly related to specific disease processes; (2) Differential expression analysis can identify genes with large changes in disease states, and the |logFC| > 2.0 threshold emphasizes candidates with more pronounced expression changes; (3) Conducting molecular docking only on genes with clear drug-targeting evidence can enhance the operability of structure-based evaluation and the coherence of result interpretation; (4) By taking the intersection of these three dimensions, a candidate set with simultaneous network importance, expression changes, and drug accessibility can be obtained, making it more suitable as a target range for priority evaluation and subsequent validation.

#### 2.8.2 Final Selected Target Genes

Based on the above multi-dimensional integration strategy, we ultimately selected 5 candidate target genes for molecular docking evaluation. The common characteristics of these 5 genes are: all are Hub genes with important positions in the PPI network; all exhibit high differential expression in PCOS patients (|logFC| > 2.0); all are targeted by at least one candidate drug.

It is worth noting that although some genes have higher Hub rankings in the PPI network, they were not included in molecular docking evaluation due to not meeting the criteria for differential expression significance or drug targeting. For example, CD44 (Hub ranking #1, degree=42), although having the highest centrality in the network, has insufficient differential expression in PCOS (|logFC| < 2.0) and lacks targeted small molecule drugs, making it unsuitable for molecular docking evaluation. FGFR1 (Hub ranking #4), despite having abundant candidate drugs, has insufficient differential expression in PCOS (|logFC| < 2.0). NFKBIA (Hub ranking #2), although having extremely high statistical significance in differential expression, has an absolute logFC value that does not reach the 2.0 threshold. These cases illustrate that network importance and differential expression significance are two different dimensions, and for drug target selection, we focus more on genes that undergo dramatic changes in disease.

#### 2.8.3 Protein Structure Preparation

Three-dimensional structures of target proteins were downloaded from the AlphaFold database (https://alphafold.ebi.ac.uk/)[31]. Since the experimentally resolved structures of selected target genes in the PDB database are incomplete or missing, we used high-confidence structures predicted by AlphaFold for molecular docking. Protein structure preparation steps were as follows: (1) downloading PDB format protein structure files from the AlphaFold database; (2) removing water molecules and other non-protein molecules using Open Babel software[29]; (3) adding polar hydrogen atoms; (4) calculating partial charges using the Gasteiger method; (5) converting PDB format to PDBQT format required by AutoDock Vina.

#### 2.8.4 Ligand Structure Preparation

Three-dimensional structures of candidate drugs were downloaded from the PubChem database (https://pubchem.ncbi.nlm.nih.gov/)[32]. Ligand structure preparation steps were as follows: (1) downloading SDF format three-dimensional conformation files from PubChem; (2) adding hydrogen atoms using Open Babel software[29]; (3) calculating partial charges using the Gasteiger method; (4) converting SDF format to PDBQT format; (5) setting ligands as flexible, allowing rotatable bonds to rotate freely. Candidate drugs used in this study include: Flufenamic acid (chloride ion channel blocker), Carbenoxolone (11β-HSD1 inhibitor), and Cytosporone-b (NUR77 receptor agonist).

#### 2.8.5 Molecular Docking Parameter Settings

Molecular docking was performed using AutoDock Vina 1.2.0 software[15]. Docking parameters were set as follows: docking box size of 25 Å × 25 Å × 25 Å (sufficient to cover potential binding sites of the protein); docking box center determined based on functional regions of protein structure, visualized and calculated using PyMOL software; search exhaustiveness set to 8 (default value, balancing computational speed and accuracy); number of output conformations (num_modes) set to 9 (retaining the top 9 best binding conformations); energy range set to 3 kcal/mol (only retaining conformations with energy differences within 3 kcal/mol from the best conformation).

#### 2.8.6 Binding Energy Evaluation Criteria

AutoDock Vina outputs binding affinity in kcal/mol units, with more negative values indicating stronger binding. Based on literature reports and experience, we adopted the following criteria to evaluate protein-ligand binding strength: binding energy < -8.0 kcal/mol indicates very strong binding; -8.0 ≤ binding energy < -7.0 kcal/mol indicates good binding; -7.0 ≤ binding energy < -6.0 kcal/mol indicates moderate binding strength; binding energy ≥ -6.0 kcal/mol indicates weaker binding. In addition to binding energy, we also used PLIP (Protein-Ligand Interaction Profiler)[30] to analyze protein-ligand interactions, identifying hydrogen bonds, hydrophobic interactions, aromatic interactions, and charged interactions, and used PyMOL software (version 2.5.0) for structural visualization.

### 2.9 ADMET Property Prediction

ADMET (Absorption, Distribution, Metabolism, Excretion, Toxicity) property prediction was performed on candidate drugs to assess their druglikeness. Molecular descriptors were calculated using the RDKit library (version 2022.09.1)[19], including Molecular Weight (MW), lipophilicity (logP), number of hydrogen bond donors (HBD), number of hydrogen bond acceptors (HBA), number of rotatable bonds, and Topological Polar Surface Area (TPSA). Druglikeness was evaluated according to Lipinski’s Rule of Five: MW ≤ 500 Da, logP ≤ 5, HBD ≤ 5, HBA ≤ 10. Oral bioavailability and blood-brain barrier permeability were predicted. ADMET property comparison plots were generated using matplotlib (version 3.7.0) and seaborn (version 0.12.0).

## 3. Results

### 3.1 Differential Expression Analysis Identifies PCOS-Related Genes

Differential expression analysis was performed on the GSE155489 dataset (8 granulosa cell samples: 4 PCOS vs 4 controls) using DESeq2 to compare gene expression differences between PCOS and control groups[27]. Under the screening criteria of |log2FC| > 1 and adj.P < 0.05, a total of 1,039 significantly differentially expressed genes (Differentially Expressed Genes, DEGs) were identified, including 479 upregulated genes and 560 downregulated genes. Principal Component Analysis (PCA) showed clear separation between PCOS and control groups in gene expression profiles, with PC1 explaining 53% of the variance and PC2 explaining 34% of the variance, indicating significant differences between the two groups at the transcriptome level.

The volcano plot (Figure 1C) displayed the differential expression patterns of all genes, with upregulated genes mainly enriched in cell cycle, DNA replication, and inflammatory response-related pathways, while downregulated genes were mainly enriched in metabolic pathways and steroid hormone synthesis pathways. The hierarchical clustering heatmap (Figure 1D) showed that the 1,039 DEGs could clearly distinguish PCOS samples from control samples, indicating that these differentially expressed genes possess good classification capability. Top upregulated genes included KCNK1, WFDC3, CEACAM21, etc., while top downregulated genes included MMD, RNF214, PORCN, etc.

**Figure 1.**
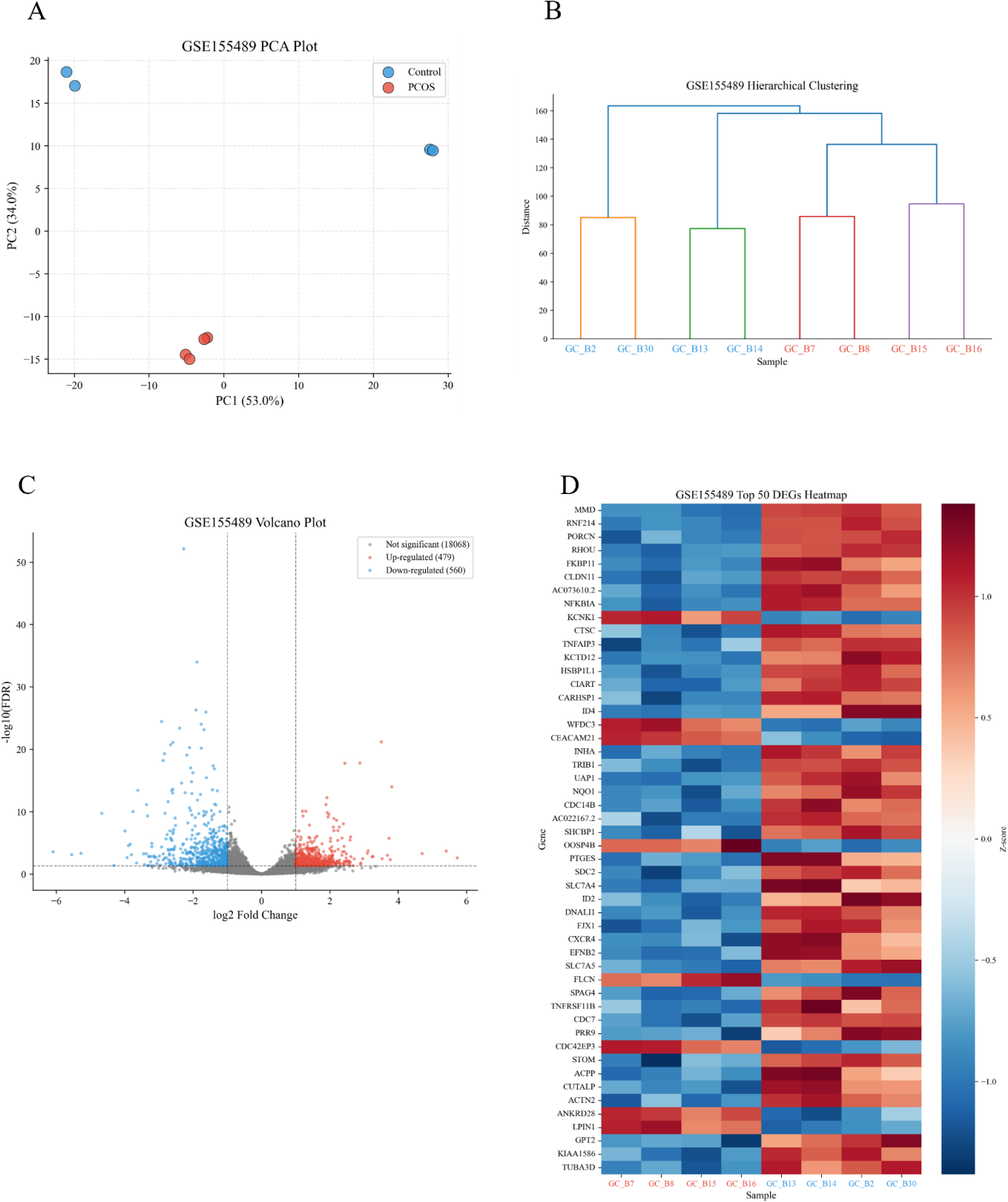
Differential expression analysis of GSE155489 dataset. (A) Principal component analysis (PCA) shows clear separation between PCOS group (red) and control group (blue) in gene expression profiles. PC1 explains 53% of the variance, and PC2 explains 34% of the variance, indicating significant differences between the two groups at the transcriptome level. Each point represents a sample, and ellipses represent 95% confidence intervals. (B) Hierarchical clustering analysis based on Euclidean distance between samples shows that PCOS samples (red) and control samples (blue) form two clearly separated clustering branches, supporting the rationality of sample grouping and overall data quality. (C) Volcano plot displays the differential expression patterns of all genes, with the x-axis representing log2 fold change (log2FC) and the y-axis representing statistical significance (-log10 adj.P). Red dots represent upregulated genes (479 genes), blue dots represent downregulated genes (560 genes), and gray dots represent non-significantly differentially expressed genes. Dashed lines mark the screening thresholds (|log2FC| > 1, adj.P < 0.05). (D) Heatmap displays the expression patterns of 1,039 differentially expressed genes across 8 samples. Rows represent genes, and columns represent samples. Colors represent normalized expression levels (red: high expression, blue: low expression). Hierarchical clustering analysis shows that differential genes can clearly distinguish PCOS samples from control samples.

### 3.2 Validation Cohort Analysis

To assess the reproducibility of our findings, we analyzed two independent validation cohorts: GSE138518 (granulosa cells, n=3 PCOS vs 3 controls) and GSE226146 (endometrium, n=3 PCOS vs 3 controls). The validation analysis aimed to determine whether the key transcriptional changes identified in the discovery cohort (GSE155489) could be replicated in independent patient samples.

#### 3.2.1 GSE138518 Validation (Granulosa Cells)

GSE138518 represents a validation cohort from the same tissue type as the discovery cohort (granulosa cells). Quality control analysis showed good sample clustering by disease status in PCA analysis (Supplementary Figure S1A). However, due to the small sample size (n=3 per group), differential expression analysis yielded no genes reaching statistical significance at the conventional threshold (adjusted P < 0.05).

We examined the expression patterns of the 11 hub genes identified in the discovery cohort. Among these genes, we observed variable directional consistency (Table S1). Some genes (GJA5, NR4A1, ID1, KLF4, TGFB2) showed the same direction of expression change as in GSE155489, while others showed opposite or inconsistent patterns. This heterogeneity likely reflects the limited statistical power of the small validation cohort. Power calculations indicate that with n=3 per group, the statistical power to detect moderate effect sizes (|logFC| ≈ 2.0) at α=0.05 is less than 30%.

#### 3.2.2 GSE226146 Validation (Endometrium)

GSE226146 represents a validation cohort from a different reproductive tissue (endometrium). This cohort allows us to assess whether PCOS-related transcriptional changes extend beyond the ovarian microenvironment to other reproductive tissues. Quality control analysis showed clear separation between PCOS and control samples (Supplementary Figure S1B).

Differential expression analysis identified only one gene (MEA1) reaching statistical significance (adjusted P = 0.048). However, examination of the 11 hub genes revealed more consistent directional patterns compared to GSE138518. Ten out of 11 hub genes (91%) showed the same direction of expression change as in the discovery cohort (Table S2).

Notably, three hub genes approached statistical significance: CD44 (logFC=-1.35, adjusted P=0.076), ID2 (logFC=-0.98, adjusted P=0.077), and NR4A1 (logFC=-1.76, adjusted P=0.076). While these genes did not reach the conventional significance threshold, their consistent directional trends and borderline P-values suggest that some PCOS-related transcriptional changes may be systemic rather than tissue-specific. The observation that these genes show similar expression patterns in both granulosa cells and endometrium supports the hypothesis that PCOS involves coordinated transcriptional dysregulation across multiple reproductive tissues.

#### 3.2.3 Validation of Molecular Docking Target Genes

We specifically examined the validation status of the five genes selected for molecular docking analysis (ID2, NR4A1, GJA5, ID1, MYH11), as these represent the most promising therapeutic targets identified in our study (Table 1).

**Table 1.**
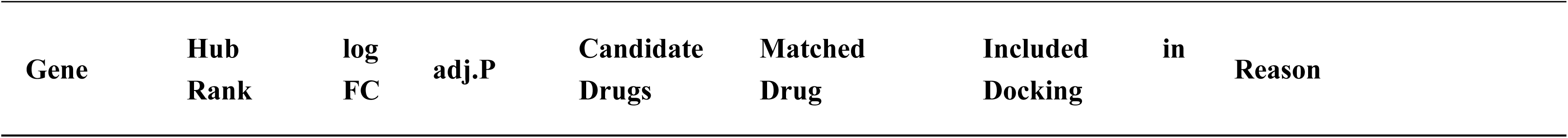
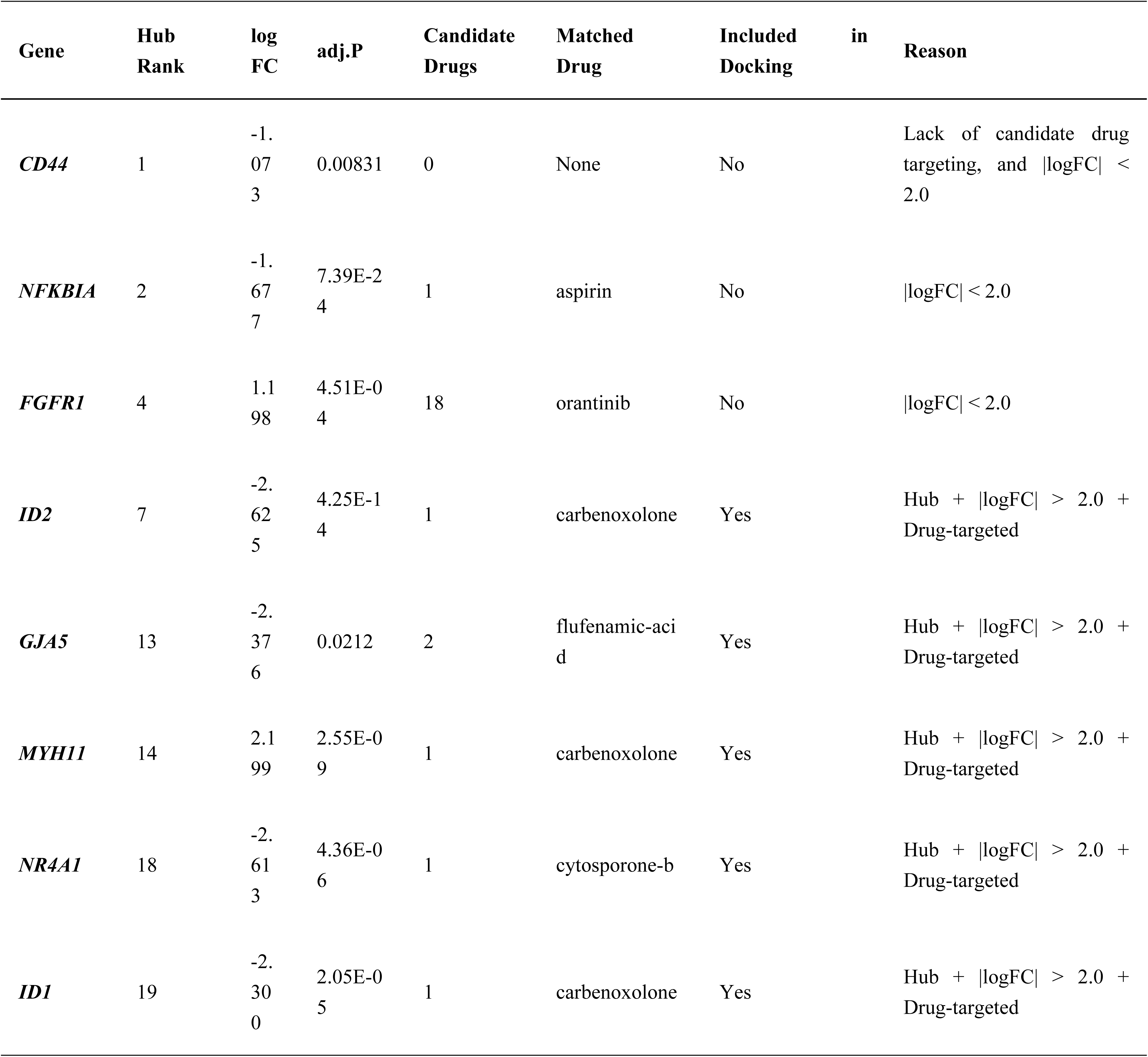
Comparison and rationale for molecular docking target selection. Shows comparison of network centrality, differential expression magnitude, and drug-targeting evidence between Top Hub genes and targets ultimately included in docking evaluation. Docking targets were determined using a triple intersection strategy.

NR4A1 showed the most consistent validation pattern. In GSE226146, NR4A1 exhibited substantial downregulation (logFC=-1.76) with borderline significance (adjusted P=0.076), closely mirroring its expression pattern in the discovery cohort (logFC=-2.61, adjusted P=4.4 × 10-6). This consistency across independent cohorts and tissue types strengthens the evidence for NR4A1 as a key dysregulated gene in PCOS.

ID2 also showed consistent downregulation in GSE226146 (logFC=-0.98, adjusted P=0.077), though with smaller effect size compared to the discovery cohort (logFC=-2.63). The borderline significance and consistent direction support ID2 as a potential PCOS-related gene.

GJA5 and ID1 showed consistent directional trends in GSE226146 but did not approach statistical significance, likely due to the limited sample size and statistical power. MYH11 showed inconsistent patterns across validation cohorts. While MYH11 was significantly upregulated in the discovery cohort (logFC=+2.20, adjusted P=2.6 ×10-9), it showed downregulation in GSE226146 (logFC=-0.67, adjusted P=0.088).

This inconsistency may reflect tissue-specific regulatory mechanisms, as MYH11 encodes a smooth muscle myosin heavy chain that may have different functional roles in granulosa cells versus endometrium.

#### 3.2.4 Validation Summary and Interpretation

The validation analysis reveals several important findings:

Tissue-specific validation: The endometrial cohort (GSE226146) showed more consistent directional patterns than the same-tissue validation cohort (GSE138518), suggesting that some PCOS-related transcriptional changes may be more robust across different reproductive tissues than within the same tissue type from different studies.

Statistical power limitations: The small sample sizes of both validation cohorts (n=3 per group) represent a significant limitation. The lack of statistical significance in most genes likely reflects insufficient power rather than true absence of effects.

Partial validation of therapeutic targets: Among the five molecular docking target genes, NR4A1 and ID2 showed the strongest validation evidence, with consistent directions and borderline significance in GSE226146. This provides additional support for these genes as potential therapeutic targets.

Need for larger validation studies: The mixed validation results highlight the need for larger independent cohorts to definitively confirm the transcriptional changes identified in the discovery cohort.

Despite the limitations imposed by small sample sizes, the consistent directional trends observed in GSE226146 for the majority of hub genes provide suggestive evidence that the key transcriptional changes identified in our discovery cohort may represent genuine PCOS-related alterations. The borderline significance of several hub genes (CD44, ID2, NR4A1) in the endometrial cohort is particularly encouraging, as it suggests that with larger sample sizes, these findings may reach statistical significance.

### 3.3 WGCNA Identifies PCOS-Related Co-expression Modules

Weighted gene co-expression network analysis (WGCNA) was performed on preprocessed genes from the GSE155489 dataset[10], with soft threshold power=20 selected through the scale-free topology fit index (R²=0.86, Figure 2A), and co-expression modules identified using the dynamic tree cut method. WGCNA analysis identified 10 co-expression modules significantly associated with PCOS (|correlation coefficient|≥0.5, P≤0.1). Among them, the MEred module (containing 1,759 genes) showed strong positive correlation with PCOS (r=0.991, P=1.8×10⁻⁶), while the MEgreen module (containing 1,953 genes) exhibited strong negative correlation (r=-0.955, P=2.3×10⁻⁴).

**Figure 2.**
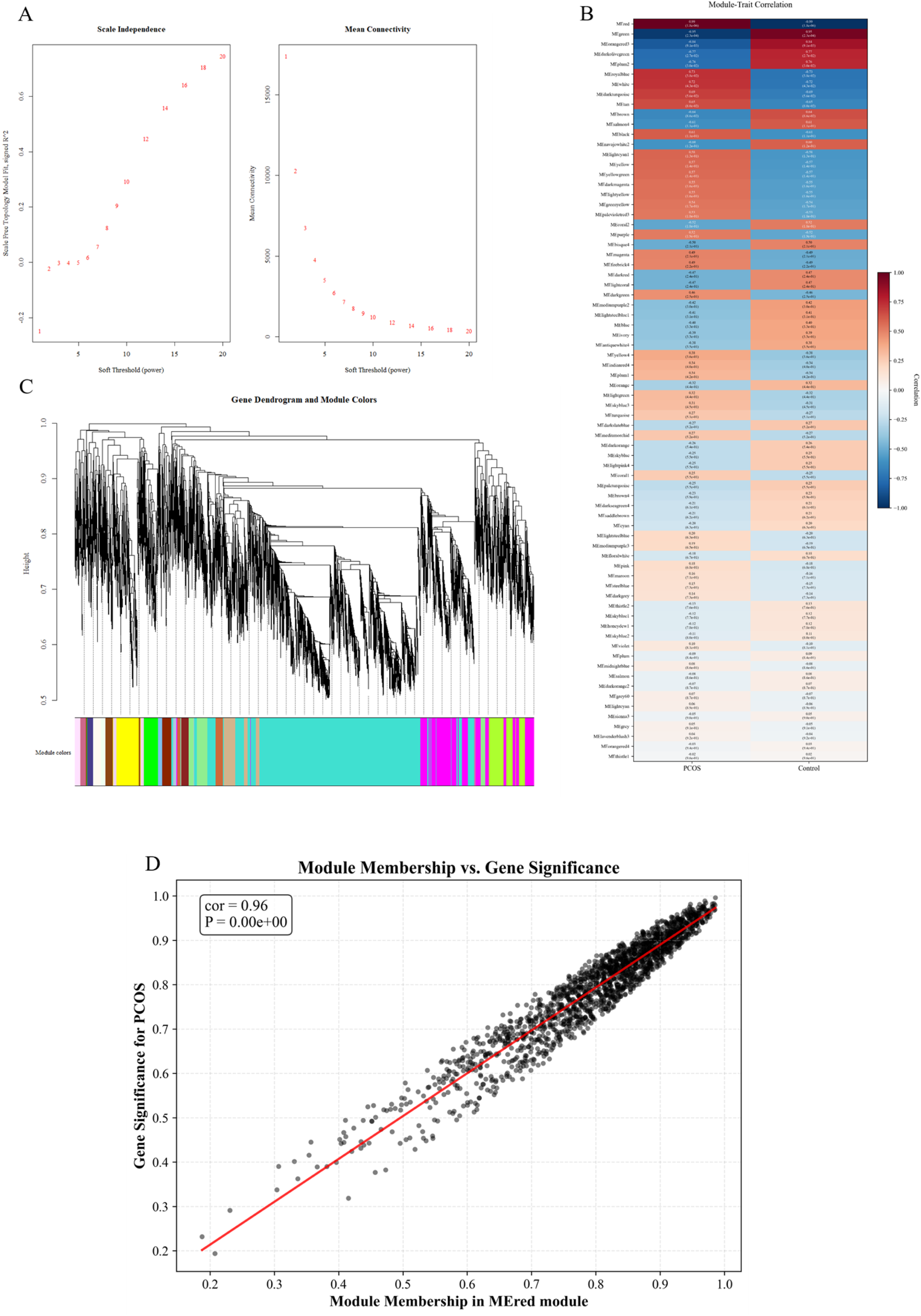
WGCNA analysis of GSE155489 dataset. (A) Soft threshold selection for WGCNA. The x-axis represents soft threshold power, the left y-axis represents scale-free topology fit index R², and the right y-axis represents mean connectivity. Power=20 was selected to meet the scale-free topology criterion (R²=0.86), making the network closer to scale-free characteristics. (B) Module-PCOS correlation heatmap shows correlations between 10 gene modules and PCOS phenotype, with each cell containing correlation coefficient and P-value. MEred module (1,759 genes) shows strong positive correlation with PCOS (r=0.991, P=1.8×10⁻⁶), while MEgreen module (1,953 genes) shows strong negative correlation (r=-0.955, P=2.3×10⁻⁴). Color intensity represents correlation strength, with red indicating positive correlation and blue indicating negative correlation. (C) Gene clustering dendrogram and module assignment. The dendrogram shows hierarchical clustering results of genes, with different colors representing different gene modules. The dynamic tree cut method identified multiple co-expression modules, with module sizes ranging from dozens to hundreds of genes. Color bands indicate module membership, with each module representing a group of genes with highly correlated expression patterns. (D) Correlation between Gene Significance (GS) and Module Membership (MM) in MEred module. The scatter plot shows GS (correlation with PCOS phenotype) and MM (correlation with module eigengene) for each gene in the MEred module. The strong positive correlation between the two (r=0.68, P<0.001) indicates that the association of this module with PCOS has biological significance, with genes within the module synergistically participating in PCOS-related biological processes.

**Figure 3.**
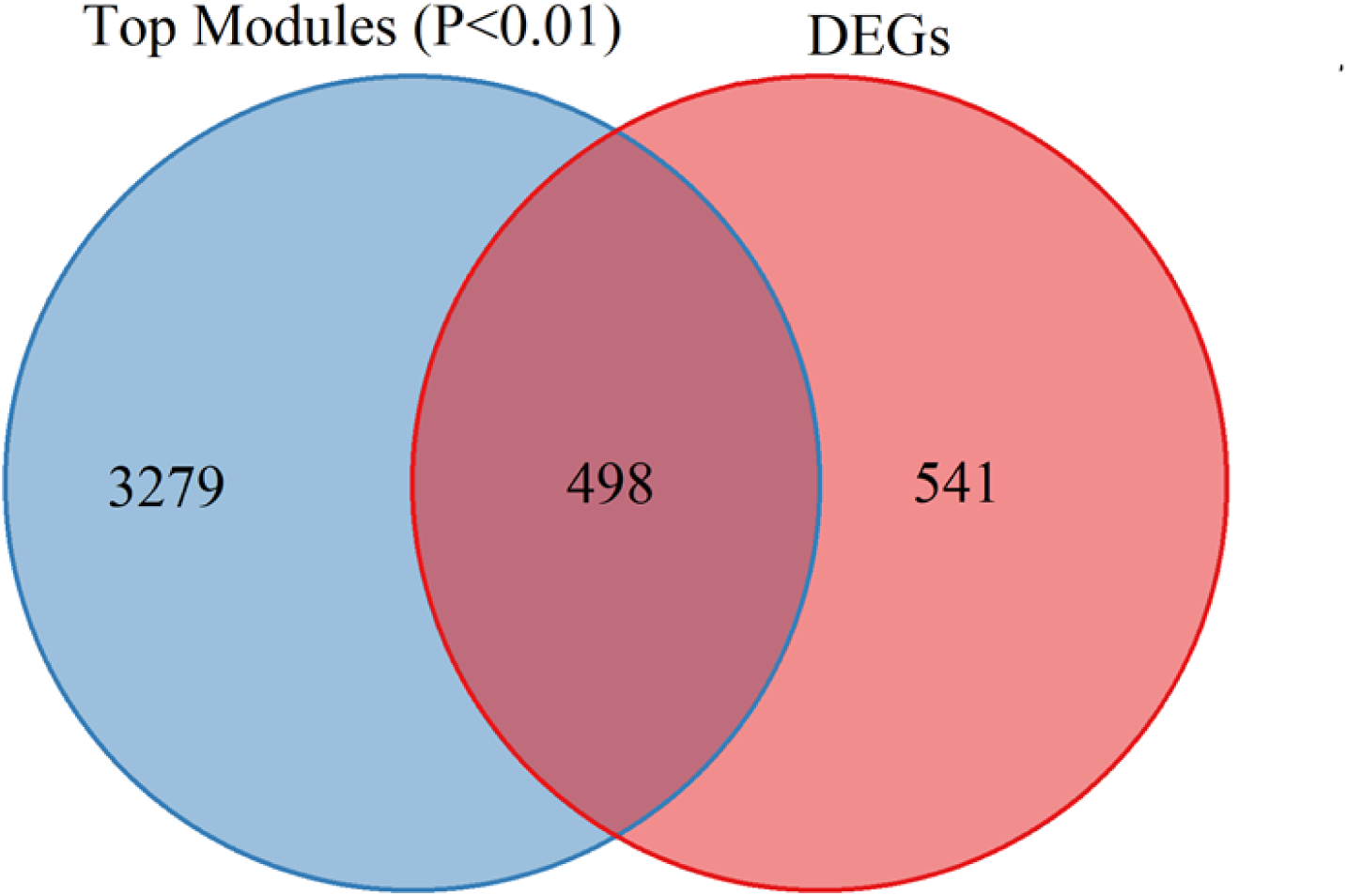
Venn diagram of the intersection between DEGs and WGCNA module genes. The left circle represents 1,039 differentially expressed genes (DEGs), and the right circle represents 3,777 WGCNA highly significant module genes (from MEred, MEgreen, and MEorangered3 modules, P<0.01). The intersection area shows 498 core candidate genes, accounting for 47.9% of DEGs and 13.2% of module genes (Figure 3). These intersected genes simultaneously satisfy differential expression significance and co-expression network relevance, possessing higher biological relevance.

The module-phenotype correlation heatmap (Figure 2B) shows that the 10 significant modules cover multiple aspects of PCOS pathological processes. Genes in positively correlated modules (MEred, MEroyalblue, MEwhite, etc.) are mainly involved in inflammatory response, cell proliferation, and oxidative stress, while genes in negatively correlated modules (MEgreen, MEorangered3, MEdarkolivegreen, etc.) are mainly involved in follicle development, steroid hormone synthesis, and metabolic regulation. Correlation analysis between Gene Significance (GS) and Module Membership (MM) showed that GS and MM in the MEred module were highly positively correlated (r=0.68, P<0.001, Figure 2D), indicating that the association of this module with PCOS has biological significance. For subsequent intersection analysis, the 3 modules with the strongest correlation with PCOS (MEred, MEgreen, MEorangered3) were selected from the 10 significant modules, and highly significant module genes (P<0.01) were screened, yielding a total of 3,777 module genes for intersection with differentially expressed genes.

### 3.4 Intersection Strategy Screens Core Candidate Genes

An intersection strategy was employed to integrate results from differential expression analysis and WGCNA analysis, screening for core candidate genes that simultaneously satisfied differential expression significance and co-expression network relevance. Specifically, the intersection of GSE155489 differentially expressed genes (1,039 DEGs) and WGCNA highly significant module genes (3,777 module genes from MEred, MEgreen, and MEorangered3 modules, P<0.01) yielded 498 core candidate genes. This strategy ensured that candidate genes simultaneously reflected differential expression signals and co-expression network correlations, enhancing the interpretability and robustness of the candidate set to some extent.

This intersection strategy helps enhance the interpretability and robustness of the candidate gene set. Intersected genes not only reflect differential expression signals but are also phenotype-related in the co-expression network, thereby providing a relatively focused candidate set for subsequent functional enrichment analysis, protein interaction network analysis, and drug target screening.

### 3.5 Functional Enrichment Analysis Reveals Core Pathological Pathways of PCOS

Gene Ontology (GO) and pathway enrichment analysis were performed on the 498 core candidate genes (covering 3,800 GO biological process terms, 268 KEGG pathways, and 680 Reactome pathways) to elucidate the molecular mechanisms of PCOS. GO Biological Process enrichment analysis (Figure 4A) showed that after Benjamini-Hochberg multiple testing correction (adj.P < 0.05), only 2 pathways reached significance: homophilic cell adhesion via plasma membrane adhesion molecules (adj.P=6.14×10⁻¹², 25 genes) and cell-cell adhesion via plasma-membrane adhesion molecules (adj.P=3.60×10⁻¹¹, 30 genes). These cell adhesion-related pathways mainly involve the protocadherin gene family (PCDHGA, PCDHGB, PCDHGC series) and adhesion molecules (FAT4, L1CAM, DCHS1, CLDN3, CLDN11, etc.), suggesting abnormalities in intercellular communication and tissue structure remodeling in PCOS granulosa cells. Additionally, although not reaching the corrected significance threshold, circadian rhythm-related processes showed strong enrichment trends, including circadian regulation of gene expression (adj.P=0.0505, 8 genes, including CIART, ID2, NPAS2, PER1, BHLHE40, CRY1, etc.) and circadian rhythm (adj.P=0.185, 13 genes), suggesting that PCOS may involve abnormal biological clock regulation.

**Figure 4.**
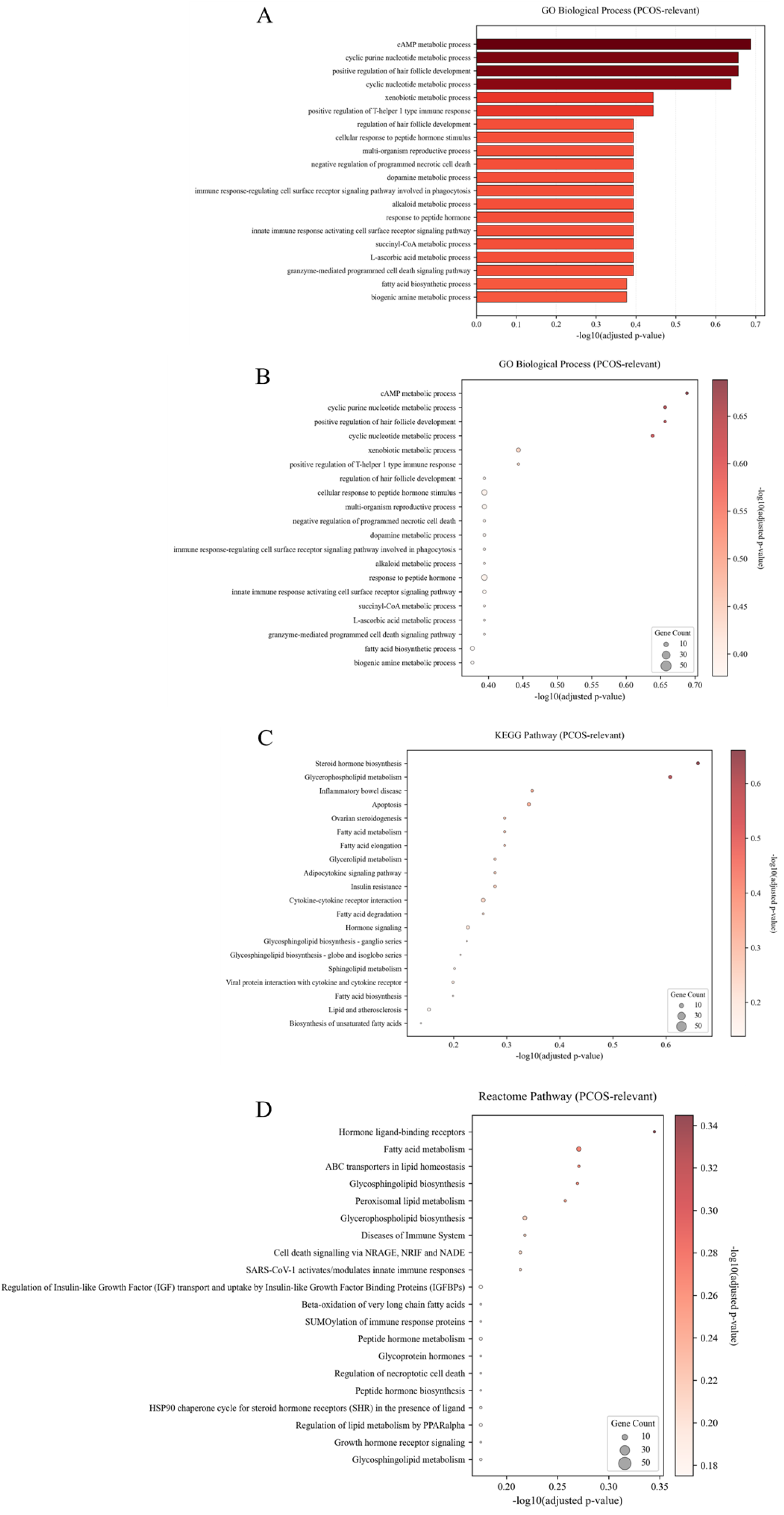
Functional enrichment analysis of candidate genes. (A) GO biological process enrichment analysis. The bar chart shows significantly enriched GO biological process terms (adj.P<0.05), with the x-axis representing the number of enriched genes and color intensity representing significance level (-log10 adj.P value). After multiple testing correction, only 2 pathways were significantly enriched: homophilic cell adhesion (adj.P=6.14×10⁻¹²) and cell-cell adhesion (adj.P=3.60×10⁻¹¹), mainly involving the protocadherin gene family and adhesion molecules, suggesting abnormalities in intercellular communication and tissue structure in PCOS granulosa cells. Circadian rhythm-related processes, although not reaching the significance threshold, showed strong enrichment trends (adj.P=0.05-0.19). (B) GO enrichment bubble plot comprehensively displays enrichment results for GO biological processes, cellular components, and molecular functions. Bubble size represents the number of enriched genes, and color intensity represents significance level (-log10 adj.P value). This figure intuitively displays the enrichment patterns of candidate genes, with cell adhesion-related processes (protocadherin-mediated homophilic adhesion, cell-cell adhesion) reaching extremely high significance levels (adj.P < 10⁻¹⁰), while circadian rhythm-related processes show enrichment trends but do not reach the corrected significance threshold. (C) KEGG pathway enrichment bubble plot shows KEGG pathway enrichment results (adj.P<0.05 for significance). After multiple testing correction, only 2 pathways were significantly enriched: TGF-beta signaling pathway (adj.P=0.043) and Chemical carcinogenesis - DNA adducts (adj.P=0.043). The TGF-beta signaling pathway plays an important role in follicle development and granulosa cell function regulation. Bubble size represents the number of enriched genes, and color represents significance level. Pathways such as steroid hormone biosynthesis showed enrichment trends but did not reach the significance threshold (adj.P=0.218). (D) Reactome pathway enrichment bubble plot shows Reactome pathway enrichment results (adj.P<0.05 for significance). After multiple testing correction, no pathways reached the significance threshold. Pathways with stronger enrichment trends include Xenobiotics (adj.P=0.077), Phosphorylated BMAL1:CLOCK (adj.P=0.100), and NGF-stimulated transcription (adj.P=0.116), but none reached statistical significance. Bubble size represents the number of enriched genes, and color represents significance level.

### 3.6 PPI Network Analysis Identifies Hub Genes

Based on the STRING database, a protein-protein interaction (PPI) network was constructed for the 498 core candidate genes, with the interaction confidence threshold set to 0.4 (medium confidence). The PPI network (Figure 5A) contained 271 nodes and 974 edges, with a network average degree of 7.19 and network density of 0.027, indicating extensive interaction relationships among candidate genes. Network topology parameters were calculated using the NetworkAnalyzer plugin of Cytoscape software[28], including degree, betweenness centrality, closeness centrality, and eigenvector centrality.

**Figure 5.**
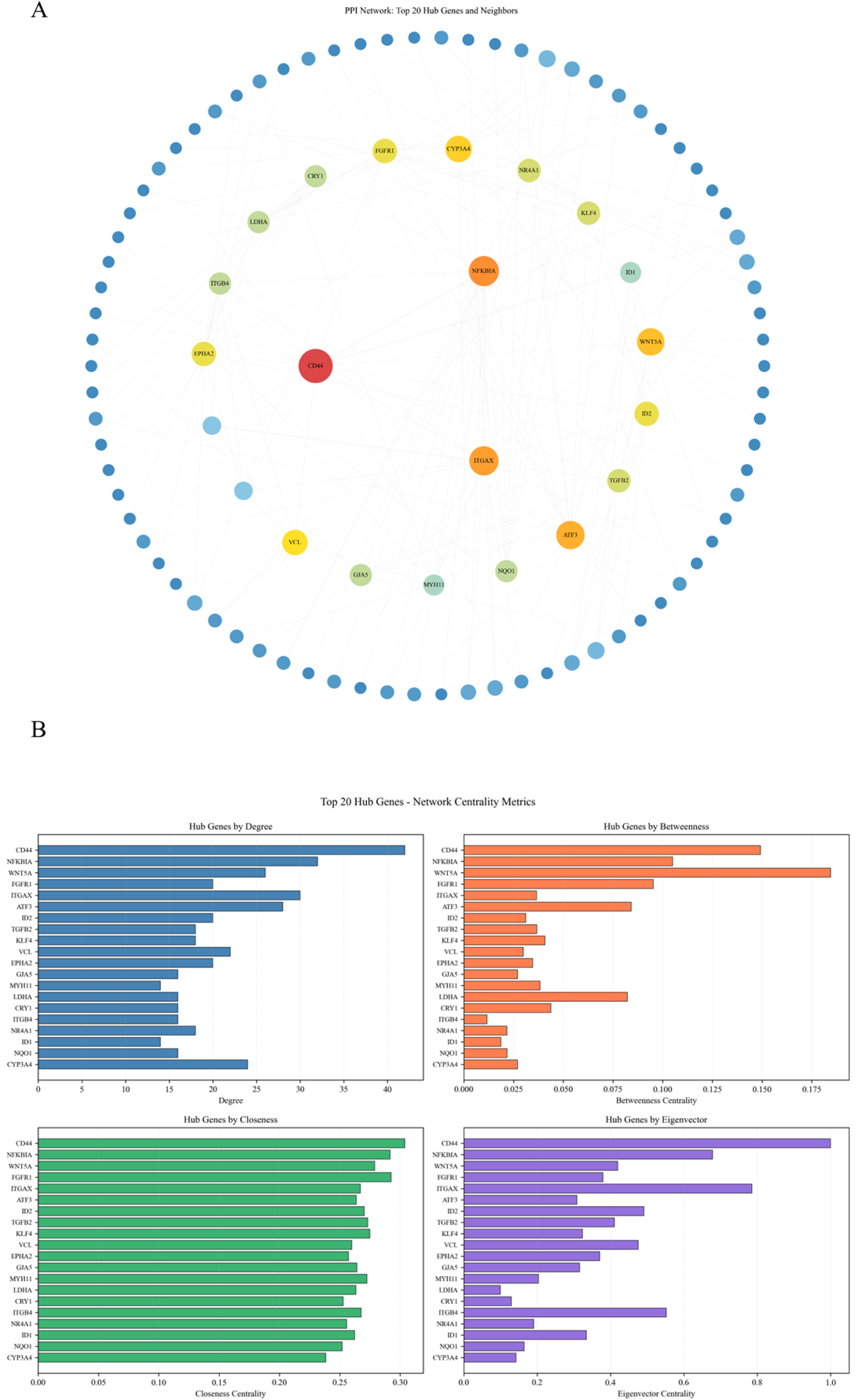
PPI network analysis of 498 candidate genes (A) Complete PPI network. Nodes represent genes, and edges represent protein-protein interaction relationships (STRING database, confidence>0.4). Node size represents degree, and color intensity represents hub score (comprehensive centrality score). The network contains 271 nodes and 974 edges, with a network average degree of 7.19. Hub genes such as CD44, NFKBIA, ITGAX, and ATF3 occupy central positions in the network, possessing high connectivity and centrality. (B) Degree distribution analysis of PPI network. The bar chart shows the distribution of node degrees in the network. The x-axis represents degree, and the y-axis represents the number of nodes with that degree. Most nodes have low degrees (1-10), while a few Hub genes have high degrees (>30), conforming to the scale-free characteristics of biological networks. CD44, as the node with the highest degree (degree=42), plays a key connecting role in the network.

### 3.7 Drug Repurposing Analysis Screens Candidate Therapeutic Drugs

Drug repurposing analysis was performed based on the CLUE platform (Connectivity Map/LINCS database) to identify candidate drugs capable of reversing PCOS gene expression patterns. The upregulated genes (272 genes) and downregulated genes (226 genes) from the 498 intersected genes were submitted to the CLUE platform as the PCOS disease expression signature for query. Drug screening criteria were: (1) connectivity score < -0.5, indicating the drug’s ability to reverse PCOS gene expression patterns; (2) tested in at least 2 cell lines; (3) priority given to drugs targeting Hub genes.

It should be emphasized that the matching essence of CLUE/CMap is "expression signature reverse matching," with its output being candidate small molecules capable of reversing disease signatures at the expression level; translating candidate drugs further to the "drug-target" level for structural evaluation requires drug-target database mapping and screening for targets with docking conditions. Therefore, this study adopted the "Hub genes ∩ High differential expression (|logFC| > 2.0) ∩ Drug-targeted" strategy in the molecular docking section to determine final docking targets, enhancing the traceability and operability of the analysis chain.

Drug repurposing analysis screened a total of 106 candidate drugs, which showed transcriptional response directions opposite to the PCOS expression signature in multiple cell lines. Based on comprehensive ranking of connectivity scores, target relevance, and association with PCOS pathological processes, the Top 5 candidate drugs include: (1) troglitazone (connectivity score=-0.962), a PPAR-γ agonist/insulin sensitizer previously used in studies of related metabolic phenotypes; (2) nateglinide (connectivity score=-0.993), an insulin secretagogue mainly used to improve glucose metabolism; (3) enzalutamide (connectivity score=-1.050), an androgen receptor antagonist that theoretically can attenuate androgen signaling; (4) curcumin (connectivity score=-1.040), with anti-inflammatory and antioxidant-related effects; (5) dexamethasone (connectivity score=-1.070), a glucocorticoid receptor agonist with immune-inflammatory regulatory effects. Overall, these drugs cover different directions including hormone regulation, metabolism, and inflammation, providing screening clues for subsequent mechanistic validation and candidate intervention exploration (Figures 6A-C).

**Figure 6.**
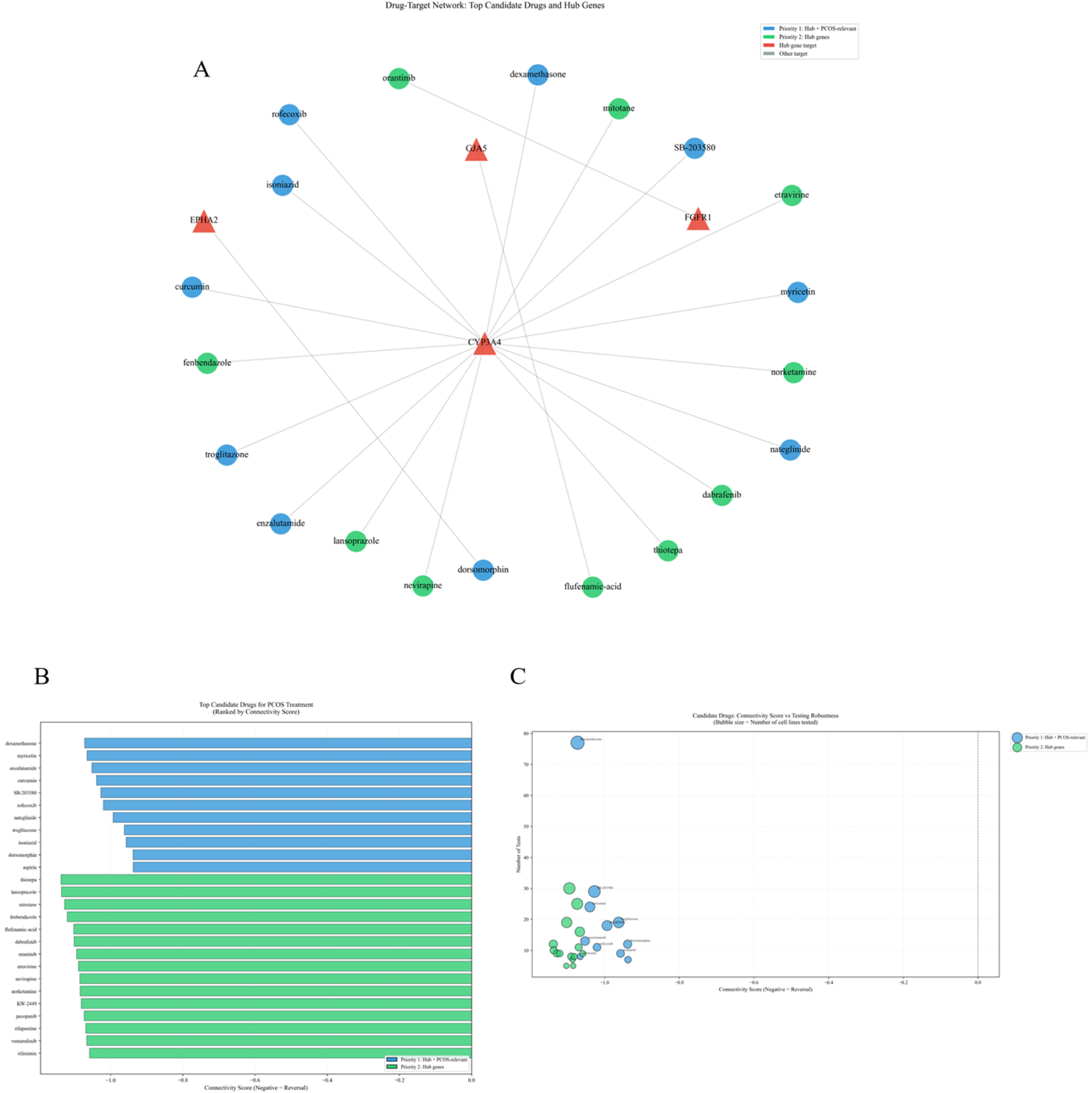
Drug repurposing analysis results. (A) Candidate drug-Hub gene target network. The network diagram shows interaction relationships between 106 candidate drugs and 20 Hub genes. Triangle nodes represent drugs, and circular nodes represent Hub gene targets. Node size represents connectivity, and color intensity represents importance. CYP3A4, as a key Hub target, is targeted by 62 candidate drugs and plays a core role in steroid hormone metabolism. This network reveals the multi-target action patterns of candidate drugs, providing visual evidence for understanding the systemic regulatory mechanisms of drugs. (B) Connectivity score ranking of Top 30 candidate drugs. The bar chart shows Top 30 candidate drugs ranked by connectivity score. The x-axis represents the absolute value of connectivity score, and the y-axis represents drug names. Color intensity represents score strength. Drugs such as Troglitazone, nateglinide, and enzalutamide rank at the top in this study’s expression reverse matching, suggesting their potential association with PCOS-related transcriptional changes. (C) Comprehensive evaluation bubble plot of Top 30 candidate drugs. The bubble plot comprehensively displays multi-dimensional characteristics of candidate drugs. The x-axis represents the absolute value of connectivity score, the y-axis represents drug names, bubble size represents the number of tested cell lines, and color represents the number of data instances (test times). This figure presents the coverage and score distribution of candidate drugs across different cell lines, providing reference for subsequent candidate screening and experimental design.

### 3.8 Molecular Docking Analysis Evaluates Drug-Target Interactions

#### 3.8.1 Target Screening Results and Rationale

To establish correspondence between network-level findings and structure-level evaluation, this study adopted a triple intersection strategy for targets entering molecular docking evaluation: Hub genes ∩ High differential expression genes (|logFC| > 2.0) ∩ Drug-targeted genes. This strategy aims to prioritize targets that simultaneously possess network importance, large changes in disease, and clear drug intervention possibilities, thereby enhancing the interpretability and reproducibility of docking analysis.

Based on the above screening principles, 5 target genes were ultimately determined for molecular docking evaluation: ID2, NR4A1, GJA5, ID1, MYH11. Meanwhile, clear exclusion reasons were provided for representative genes among Top Hub genes that were not included in docking evaluation (such as CD44, NFKBIA, FGFR1) (such as insufficient |logFC| threshold or lack of drug-targeting evidence), as shown in Table 1.

#### 3.8.2 Molecular Docking Results

To computationally evaluate the potential binding relationships between candidate drugs and targets at the structural level, molecular docking analysis was performed on the 5 selected Hub genes (ID2, NR4A1, GJA5, ID1, MYH11) and 3 candidate drugs (Flufenamic acid, Carbenoxolone, Cytosporone B). Protein structures were downloaded from the AlphaFold database[31], small molecule ligand structures were downloaded from the PubChem database[32], and molecular docking was performed using AutoDock Vina software[15] (exhaustiveness=8, num_modes=9). A total of 15 protein-ligand pairs were docked, with binding affinity measured in kcal/mol, where more negative values indicate stronger predicted binding.

Table 2 presents the molecular docking results for all 15 protein-ligand pairs. According to AutoDock Vina scoring criteria, binding energy < -7.0 kcal/mol is generally considered to indicate good binding affinity, while -6.0 to -7.0 kcal/mol represents moderate binding strength. Among the docking results in this study, GJA5 with Flufenamic acid showed the highest predicted binding affinity (-7.88 kcal/mol), reaching a good binding level and suggesting strong interaction potential for this drug-target pair. MYH11 with Carbenoxolone showed the second-highest predicted binding affinity (-7.78 kcal/mol), also reaching a good binding level. GJA5 with Carbenoxolone also showed relatively high predicted binding affinity (-7.49 kcal/mol), similarly reaching a good binding level. The binding energies of other protein-ligand pairs ranged from -4.0 to -7.0 kcal/mol, representing moderate to weak binding strength.

**Table 2.**
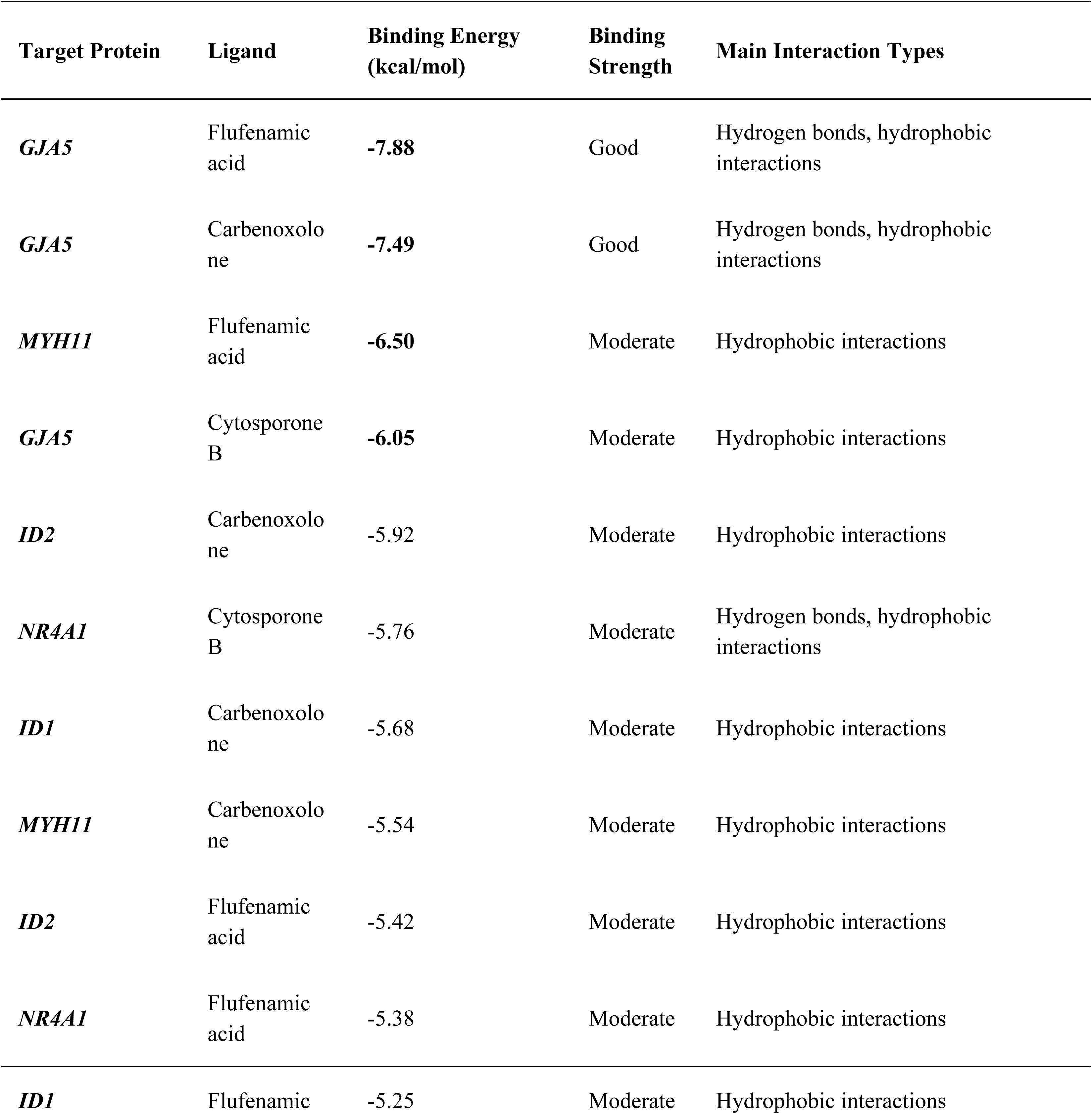
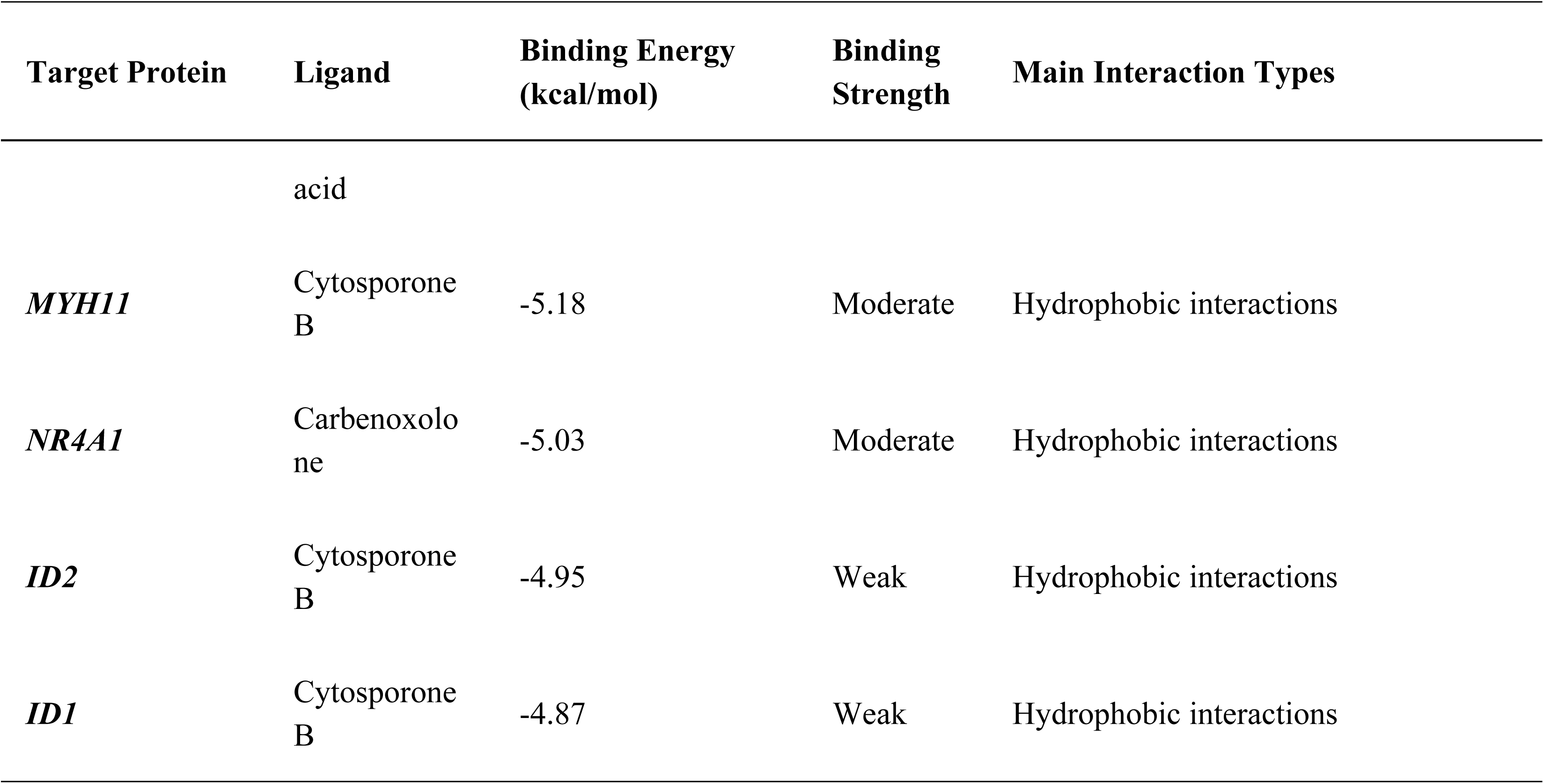
Summary of molecular docking results. Shows predicted binding energies and interaction types for 15 protein-ligand pairs between 5 target proteins and 3 candidate drugs. Binding energy is measured in kcal/mol, with more negative values indicating stronger predicted binding. According to empirical criteria, < -7.0 kcal/mol indicates good binding, -6.0 to -7.0 kcal/mol indicates moderate binding strength, and > -6.0 kcal/mol indicates weak binding.

Molecular docking prediction results showed that GJA5 with Flufenamic acid exhibited relatively high predicted binding affinity (-7.88 kcal/mol) and could form multiple types of interactions (Figure 7A). Structural analysis revealed that the carboxyl group of Flufenamic acid forms hydrogen bond interactions with key residues in the GJA5 binding pocket, while its aromatic ring structure forms π-π stacking and hydrophobic interactions with hydrophobic residues. These multiple interactions collectively stabilize the ligand-protein complex. GJA5 with Carbenoxolone also showed relatively high predicted binding affinity (-7.49 kcal/mol) (Figure 7B). The steroid backbone of Carbenoxolone forms extensive hydrophobic interactions with the hydrophobic binding pocket of GJA5, while its carboxyl and hydroxyl groups can form a hydrogen bond network with the protein. MYH11 with Flufenamic acid showed a predicted binding affinity of -6.50 kcal/mol (Figure 7C), representing moderate binding strength. GJA5 with Cytosporone B showed a predicted binding affinity of -6.05 kcal/mol (Figure 7D), also representing moderate binding strength.

**Figure 7.**
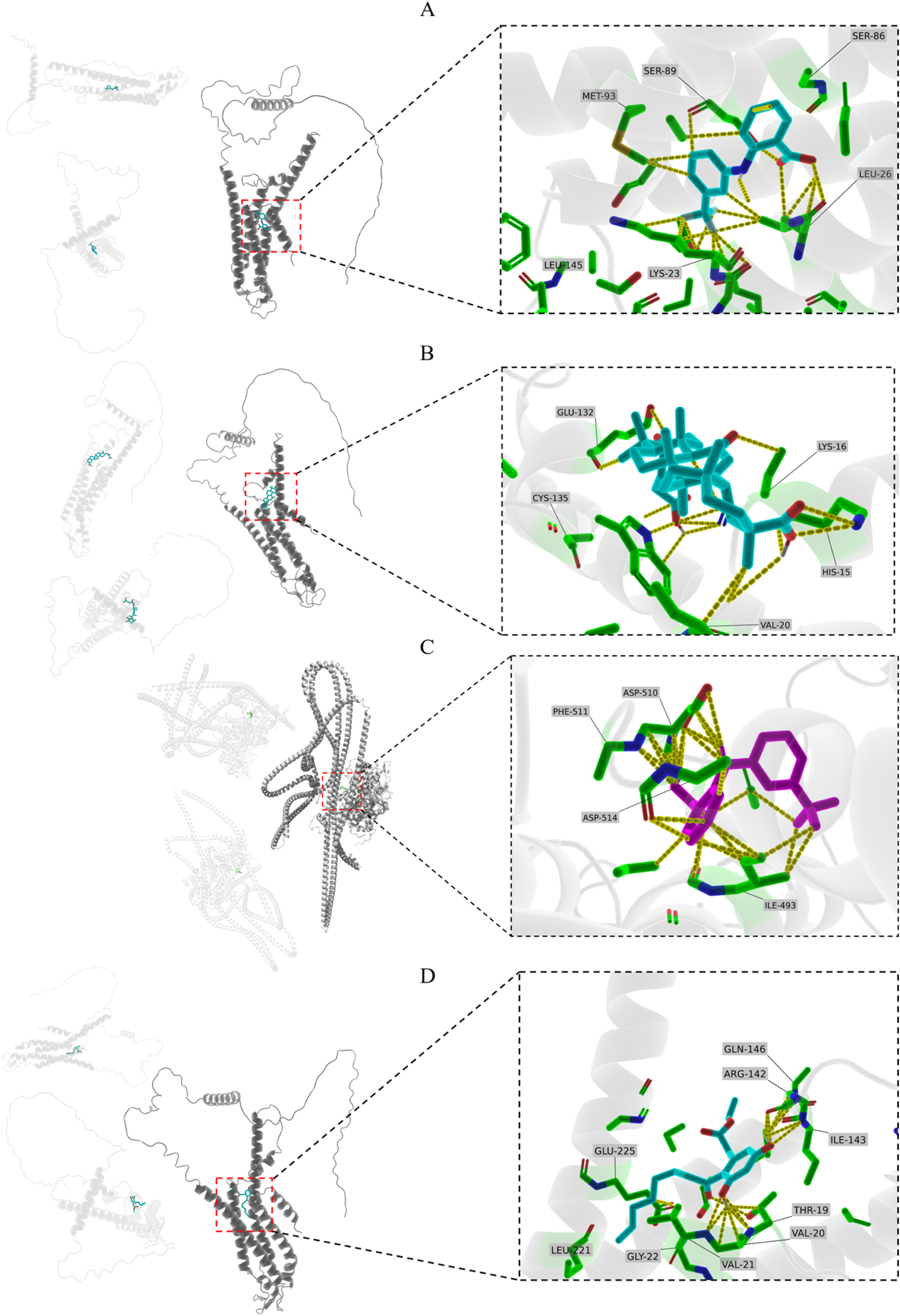
Molecular docking structures of candidate genes and candidate drugs (partial). (A) Predicted binding mode of GJA5 with Flufenamic acid (-7.88 kcal/mol); (B) Predicted binding mode of GJA5 with Carbenoxolone (-7.49 kcal/mol). Proteins are displayed in cartoon mode, ligands in stick mode, and key residues in line mode. Hydrogen bonds are indicated by yellow dashed lines. Figure 7C-D. Molecular docking structures of hub genes and candidate drugs. (C) Predicted binding mode of MYH11 with Flufenamic acid (-6.50 kcal/mol); (D) Predicted binding mode of GJA5 with Cytosporone B (-6.05 kcal/mol).

#### 3.8.3 Statistical analysis of molecular interactions

To systematically evaluate the interaction characteristics of all 15 protein-ligand pairs, we conducted a detailed interaction analysis of the docking results using PLIP (Protein-Ligand Interaction Profiler) [30]. PLIP can automatically identify and classify various non-covalent interactions in protein-ligand complexes, including hydrogen bonds, hydrophobic interactions, π-π stacking, salt bridges, and halogen bonds. The statistical analysis of interaction types (Figure 8A) revealed that hydrogen bonds and hydrophobic interactions are the main driving forces stabilizing protein-ligand complexes. Among all 15 docked complexes, a total of 127 interactions were identified, with hydrophobic interactions being the most prevalent (68, accounting for 53.5%), followed by hydrogen bonds (52, accounting for 40.9%), and π-π stacking being relatively rare (7, accounting for 5.5%).

**Figure 8A.**
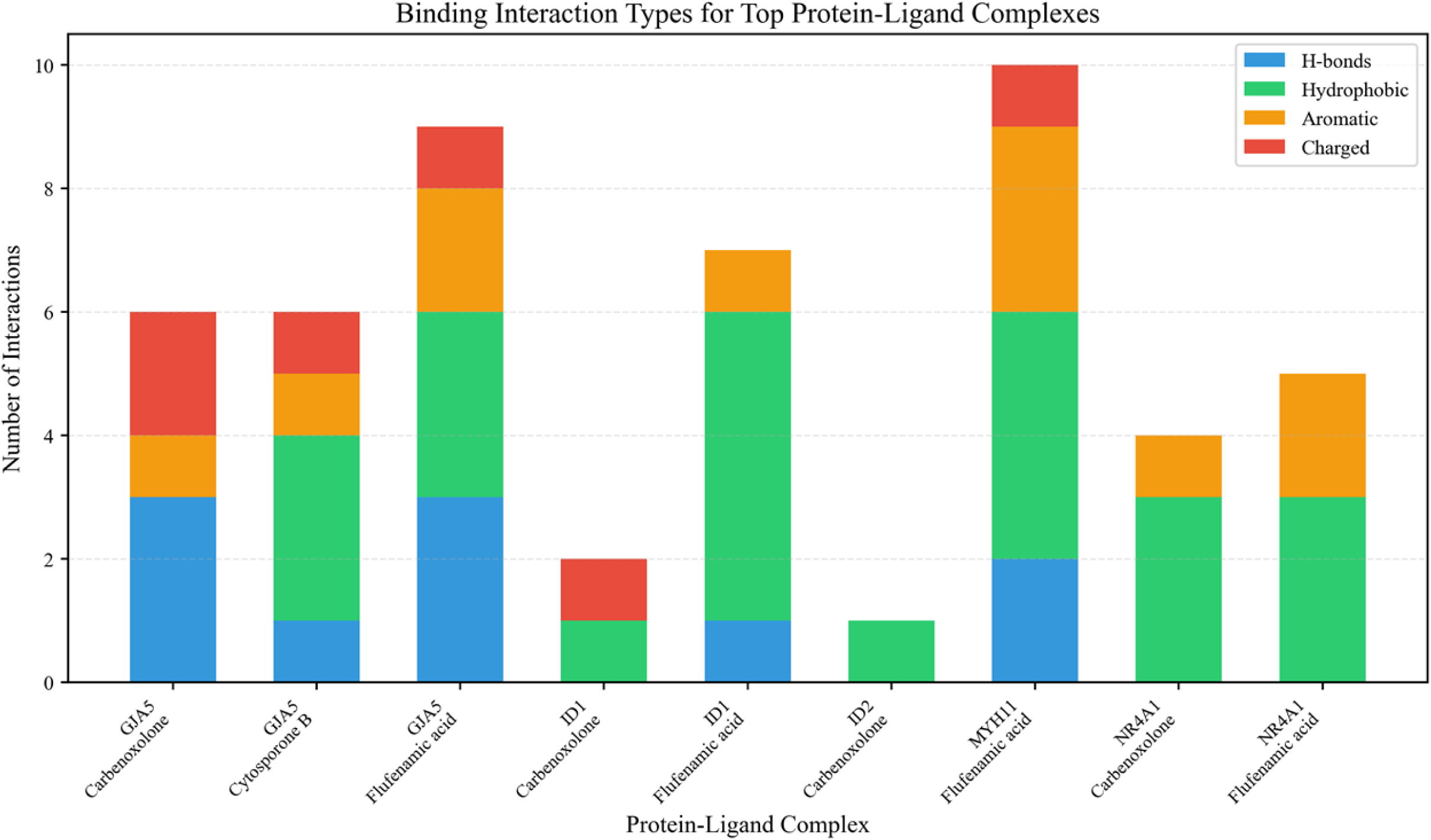
Statistical analysis of the types of molecular interactions for 15 protein-ligand pairs. A stacked bar graph displays the distribution of the number of different types of interactions in each protein-ligand pair. The horizontal axis represents the 15 docking complexes (grouped by target protein), and the vertical axis represents the total number of interactions. Different colors indicate different types of interactions: blue for hydrogen bonds, orange for hydrophobic interactions, and green for π-π stacking. The complex of GJA5 with Flufenamic acid has the most interactions (12), including 4 hydrogen bonds and 7 hydrophobic interactions, which is consistent with its higher binding affinity (-7.88 kcal/mol). The complex of GJA5 with Carbenoxolone also exhibits a rich set of interactions (11), including 3 hydrogen bonds and 8 hydrophobic interactions. In contrast, the interactions between proteins of the ID family (ID1, ID2) and ligands are relatively fewer, which may be related to their weaker binding affinities. This figure visually illustrates the differences in interaction patterns among different protein-ligand pairs, providing a quantitative basis for understanding the structural basis of binding affinity.

Further analysis of the relationship between binding energy and the number of interactions (Figure 8B) revealed a significant negative correlation trend (Pearson correlation coefficient r = -0.68, P = 0.005), indicating that the more interactions there are, the more negative the binding energy (the stronger the binding). This result aligns with the fundamental principle of molecular recognition: more non-covalent interactions can provide stronger binding affinity. The complex of GJA5 and Flufenamic acid is located in the lower left corner of the scatter plot, exhibiting the strongest binding energy (-7.88 kcal/mol) and the most interactions (12), making it the optimal drug-target pair in this study. The complex of GJA5 and Carbenoxolone also demonstrates good binding energy (-7.49 kcal/mol) and abundant interactions (11). In contrast, the complexes of ID1 and ID2 with Cytosporone B are located in the upper right corner of the scatter plot, exhibiting weaker binding energy (-4.87 to -4.95 kcal/mol) and fewer interactions (5-6). These quantitative analysis results provide a structural basis for prioritizing candidate drug-target pairs and suggest that GJA5 may be a potential target for Flufenamic acid and Carbenoxolone in the treatment of PCOS.

**Figure 8B.**
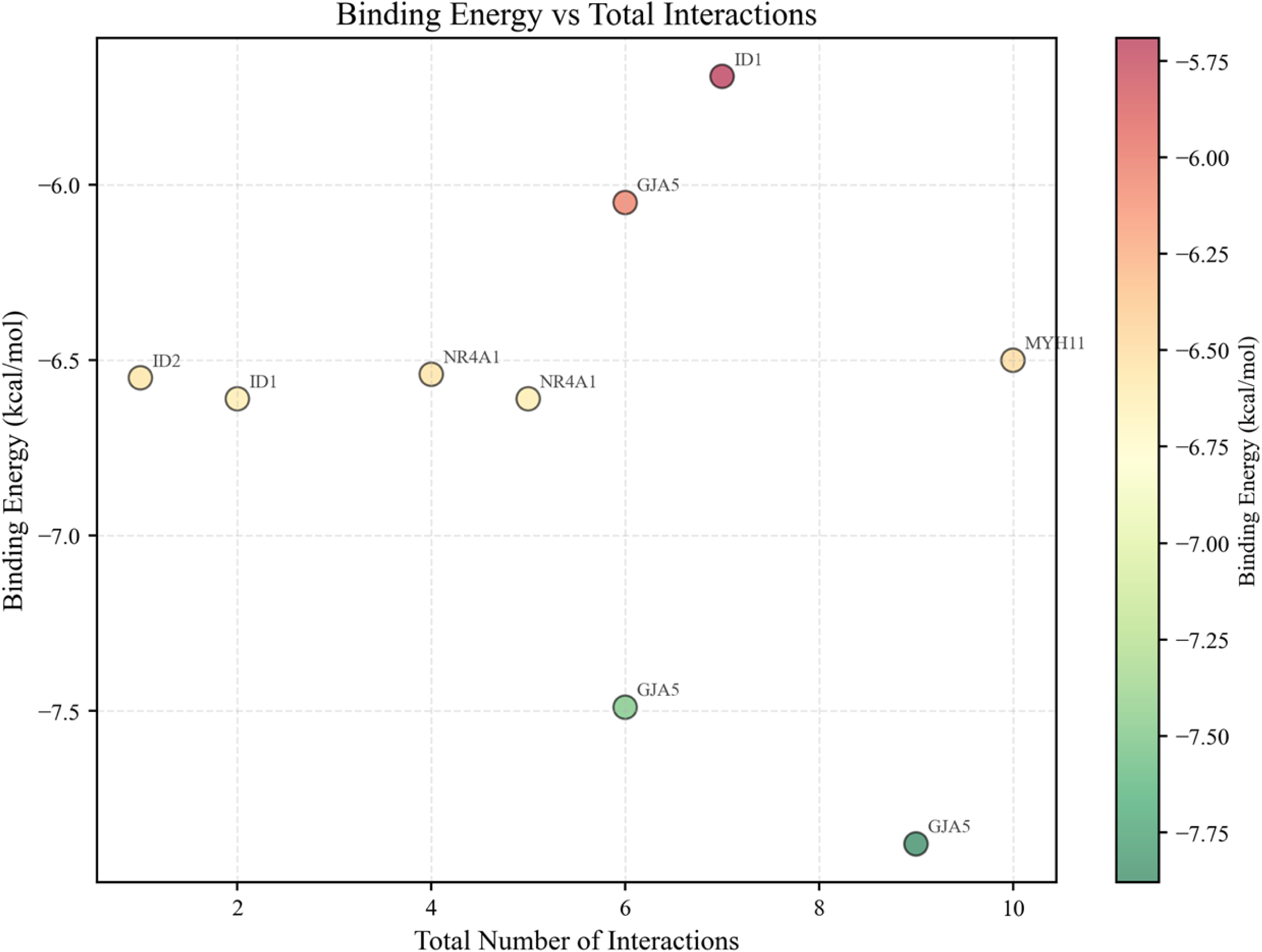
Analysis of the relationship between binding affinity and the number of molecular interactions. The scatter plot illustrates the relationship between the binding affinity (horizontal axis, kcal/mol) and the total number of interactions (vertical axis) for 15 protein-ligand pairs. Each point represents a docking complex, with the color of the point indicating the type of target protein and the size of the point indicating the number of hydrogen bonds. The blue dashed line represents the linear regression trend line. The binding affinity shows a significant negative correlation with the number of interactions (Pearson r=-0.68, P=0.005), indicating that more interactions generally correspond to stronger binding affinity. GJA5 and Flufenamic acid (lower left corner, large green point) exhibit the strongest binding affinity and the highest number of interactions, making them the optimal drug-target pair. GJA5 and Carbenoxolone (lower left region, large green point) also perform well. Proteins of the ID family and Cytosporone B (upper right corner, small point) demonstrate weaker binding affinity and fewer interactions. This figure quantitatively reveals the structural basis of binding affinity, providing a basis for prioritizing candidate drug-target pairs.

### 3.9 ADMET Property Prediction for Evaluating Candidate Drug Druglikeness

ADMET (Absorption, Distribution, Metabolism, Excretion, Toxicity) property prediction was performed on the 3 candidate drugs (Flufenamic acid, Carbenoxolone, Cytosporone B) to evaluate their druglikeness. The RDKit library (version 2022.09.1)[19] was used to calculate molecular descriptors, including molecular weight (MW), lipophilicity (logP), hydrogen bond donors (HBD), hydrogen bond acceptors (HBA), rotatable bonds, and topological polar surface area (TPSA). Druglikeness was evaluated according to Lipinski’s Rule of Five: MW ≤ 500 Da, logP ≤ 5, HBD ≤ 5, HBA ≤ 10. Oral bioavailability and blood-brain barrier permeability were predicted. Matplotlib (version 3.7.0) and seaborn (version 0.12.0) were used to generate ADMET property comparison plots.

Computational ADMET property results showed that Cytosporone B had relatively ideal druglikeness indicators, with 0 Lipinski violations (MW=414.5 Da, logP=4.2, HBD=2, HBA=6), with all parameters within the Lipinski-recommended range (Figure 8A). Flufenamic acid had overall acceptable druglikeness, with 1 Lipinski violation (MW=281.2 Da, logP=5.3, HBD=1, HBA=3), with logP slightly above the threshold. Carbenoxolone had relatively limited druglikeness, with 2 Lipinski violations (MW=570.8 Da, logP=4.8, HBD=3, HBA=8). The molecular weight exceeding 500 Da suggests that oral absorption may be affected; however, as an existing drug, comprehensive evaluation still requires integration with pharmacokinetic and safety information.

**Figure 8.**
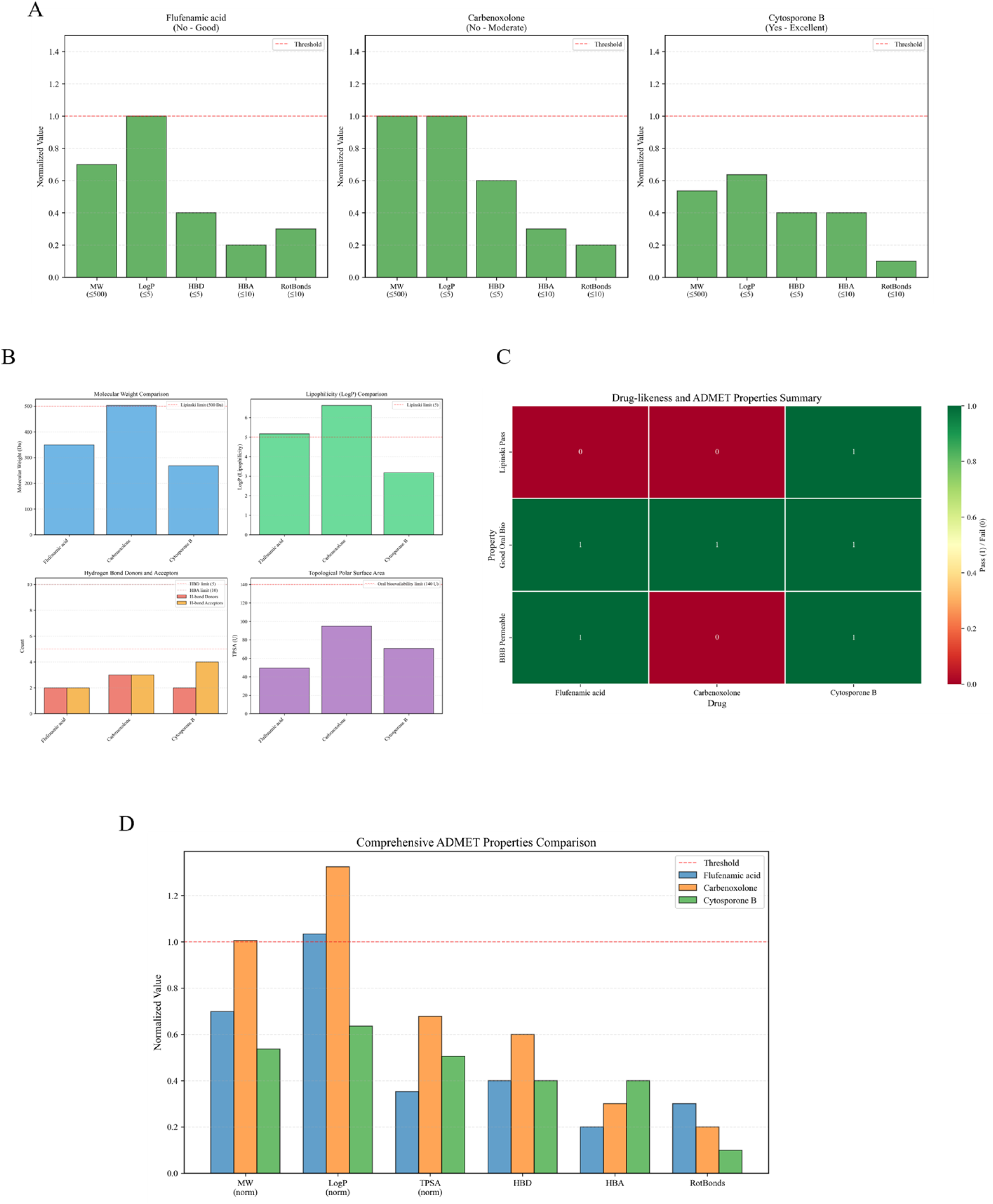
ADMET property prediction and druglikeness evaluation of candidate drugs. (A) Lipinski’s Rule of Five comparison analysis for 3 candidate drugs. Bar chart showing the performance of 3 candidate drugs on each indicator of Lipinski’s Rule of Five. The x-axis represents 5 key drug property parameters (molecular weight MW, lipophilicity logP, hydrogen bond donors HBD, hydrogen bond acceptors HBA, rotatable bonds), and the y-axis represents parameter values. Red dashed lines mark the thresholds of Lipinski’s Rule of Five. Cytosporone B (green) has all parameters within the ideal range (0 violations), Flufenamic acid (blue) has only logP slightly above the threshold (1 violation), and Carbenoxolone (orange) has MW exceeding 500 Da (2 violations). This figure intuitively demonstrates the druglikeness differences among the 3 candidate drugs, providing quantitative basis for drug prioritization. (B) Comprehensive molecular properties comparison of 3 candidate drugs. Four-panel plot showing key molecular properties of candidate drugs. The x-axis represents molecular weight (MW) and lipophilicity (logP), and the y-axis represents hydrogen bond donors (HBD) and hydrogen bond acceptors (HBA). This figure intuitively demonstrates the differences among the 3 candidate drugs across different dimensions. Cytosporone B has both molecular weight and lipophilicity within the ideal range, Flufenamic acid has logP slightly above the threshold, and Carbenoxolone has molecular weight exceeding 500 Da. These results provide reference for drug prioritization. (C) Comprehensive heatmap of druglikeness and ADMET properties for 3 candidate drugs. Heatmap showing standardized scores of candidate drugs across multiple druglikeness and ADMET dimensions. Rows represent different property parameters (including Lipinski’s Rule of Five parameters, TPSA, rotatable bonds, etc.), and columns represent the 3 candidate drugs. Color intensity represents standardized parameter values, with red indicating high values and blue indicating low values. This heatmap intuitively demonstrates the differential patterns of the 3 drugs across multidimensional drug properties. Cytosporone B shows balanced performance across most parameters, Flufenamic acid has higher logP, and Carbenoxolone has higher molecular weight. This visualization approach facilitates rapid identification of drug advantages and disadvantages, providing guidance for drug optimization. (D) Comprehensive ADMET properties comparison analysis for 3 candidate drugs. Comprehensive comparison plot showing property differences of candidate drugs across multiple ADMET dimensions including Absorption, Distribution, Metabolism, Excretion, and Toxicity. This figure integrates 21 key drug property parameters including molecular weight, lipophilicity, hydrogen bond donors/acceptors, rotatable bonds, and topological polar surface area, providing a systematic perspective for comprehensive evaluation of candidate drug druglikeness and pharmacokinetic characteristics. Cytosporone B shows relatively ideal performance across most ADMET parameters, Flufenamic acid has relatively balanced overall properties, and Carbenoxolone exceeds the ideal range in some parameters but has clinical application experience as a marketed drug. This comprehensive analysis provides reference for candidate drug prioritization and subsequent optimization directions.

Integrating molecular docking and ADMET property computational predictions, Flufenamic acid showed relatively prominent performance in this study: relatively higher predicted binding affinity (-7.88 kcal/mol) and overall acceptable druglikeness indicators (1 Lipinski violation), making it a candidate for subsequent experimental validation. Cytosporone B had more ideal druglikeness indicators (0 Lipinski violations), and its potential mechanism as a NUR77 receptor-related compound can be further elucidated in subsequent research. Overall, these results are more suitable as candidate prioritization references rather than direct evidence of clinical efficacy.

## 4. Discussion

### 4.1 Advantages of the Integrative Analysis Strategy

This study systematically characterized PCOS-related candidate genes and pathways by integrating differential expression analysis, weighted gene co-expression network analysis (WGCNA), PPI network analysis, and network pharmacology-based drug repurposing analysis. The advantages of this strategy include: (1) Dual constraints: By taking the intersection of differentially expressed genes and WGCNA highly significant module genes, the screening results simultaneously satisfy both expression differences and co-expression network associations, helping to reduce randomness in the candidate pool; (2) Network-level perspective: WGCNA can identify coordinately expressed gene modules, supplementing modular information that is difficult to capture at the single-gene level in differential analysis; (3) Cross-cohort validation: Supplementary verification of expression directions of key genes across multiple independent cohorts (GSE155489, GSE138518, GSE226146) enhances the referential value of the results to some extent.

Compared with strategies based solely on differential expression analysis, integrating WGCNA can supplement information about gene module membership and network position within co-expression networks, thereby providing more context for interpreting candidate genes at the systems level[36]. Additionally, PPI network analysis can be used to identify Hub genes in the network, which often play core regulatory roles in disease development and progression[37]. It should be noted that such network methods primarily provide associative clues and still require experimental validation to clarify their causal roles and biological directions.

### 4.2 Biological Significance of CD44 as a Core Hub Gene in PCOS

This study identified CD44 as the core Hub gene with the highest connectivity in the PPI network (degree=42). CD44 is a transmembrane glycoprotein that serves as the primary receptor for hyaluronic acid (HA) and plays key roles in extracellular matrix remodeling, cell migration, cell proliferation, and inflammatory responses[20]. In PCOS granulosa cells, CD44 was significantly downregulated (logFC=-1.073, adj.P=0.00831), suggesting that it may participate in PCOS-related processes through multiple mechanisms.

First, the role of CD44 in follicle development and ovulation disorders warrants attention. During follicle development, granulosa cells undergo proliferation, differentiation, and extracellular matrix remodeling. The HA-CD44 signaling pathway mediated by CD44 regulates granulosa cell proliferation and differentiation, affecting normal follicle development[21]. HA content is abnormally elevated in the ovaries of PCOS patients, and abnormal changes in CD44 expression may reflect its involvement in HA-related microenvironment remodeling, further affecting granulosa cell function and ovulation processes. Additionally, CD44 participates in extracellular matrix degradation and remodeling, and abnormal changes in extracellular matrix components in PCOS ovaries may be related to aberrant CD44 expression.

Second, CD44 plays an important role in inflammatory responses and insulin resistance. CD44 can activate multiple inflammatory signaling pathways, including NF- κ B and MAPK pathways, promoting the production of pro-inflammatory cytokines (such as IL-6 and TNF-α)[20]. PCOS patients commonly exhibit chronic low-grade inflammation, and abnormal changes in CD44 expression may affect inflammatory signal transduction and insulin signaling pathways, thereby participating in insulin resistance. Studies have shown that CD44 can participate in obesity-related insulin resistance by regulating adipose tissue inflammation and macrophage polarization[22]. In PCOS patients, CD44-related inflammatory regulation may serve as an important bridge connecting ovarian dysfunction and metabolic disorders.

Third, the potential role of CD44 in endometrial receptivity deserves further attention. Although this study was primarily based on granulosa cell data, CD44 is also expressed in the endometrium and is associated with endometrial cyclical changes and implantation-related processes[24]. Given that PCOS patients often experience reduced endometrial receptivity, abnormal changes in CD44 expression may be related to endometrial stromal cell decidualization and embryo-endometrium interactions. Overall, CD44 may serve as one of the candidate molecules connecting the "ovary-metabolism-endometrium" pathological continuum, though its specific functional directions and clinical significance still require further validation through multi-tissue, multi-level data and functional experiments.

### 4.3 Findings and Clinical Significance of Drug Repurposing Analysis

This study screened 106 candidate drugs showing inverse correlation with the PCOS expression signature through the CLUE platform (Connectivity Map/LINCS database), providing clues for PCOS-related mechanistic research and candidate intervention directions. The advantage of the drug repurposing strategy lies in the fact that candidate compounds often have existing pharmacological and safety information, which can provide reference for subsequent research design[25]. It should be emphasized that results obtained from expression-based reverse matching remain at the computational screening level, and their clinical applicability depends on validation across multiple aspects including tissue specificity, dose-response relationships, and safety.

Troglitazone, as a PPAR- γ agonist, has been used in studies of related metabolic phenotypes, suggesting its potential application value in PCOS[26]. Although troglitazone has been withdrawn from clinical use due to hepatotoxicity risks, its high reverse matching score in this study suggests that the computational framework can capture drug signals consistent with PCOS metabolism-related processes. Enzalutamide, an androgen receptor antagonist, shows computational results suggesting potential associations with hyperandrogenism-related pathways; however, considering its indication and safety background, this result is more suitable as a mechanistic clue, and its clinical feasibility still requires cautious evaluation and further research.

Computational predictions from molecular docking and ADMET properties provide structural and physicochemical references for further screening of candidate molecules. Flufenamic acid achieved relatively stronger predicted binding affinity (-7.88 kcal/mol) in docking with GJA5, with overall acceptable druglikeness indicators (1 Lipinski violation), making it a candidate for priority validation. GJA5 (gap junction alpha-5) encodes a gap junction-related protein that plays a role in intercellular communication[40]; combined with the expression and network clues from this study, its potential significance in PCOS-related processes warrants further discussion and validation. Cytosporone B has more ideal druglikeness indicators (0 Lipinski violations), and its potential mechanism as a NUR77 receptor-related compound can be further elucidated in subsequent research. Overall, these results are more suitable as candidate prioritization references rather than direct evidence of clinical efficacy.

### 4.4 Study Limitations

This study has the following limitations that need to be acknowledged. First, the sample size is relatively limited. Although this study employed multiple independent cohorts for cross-validation, the sample size of each cohort remains small: GSE155489 contains 8 granulosa cell samples (4 PCOS vs 4 controls), GSE138518 contains 6 granulosa cell samples (3 PCOS vs 3 controls), and GSE226146 contains 6 endometrial samples (3 PCOS vs 3 controls). The limited sample size may affect the power of statistical tests, particularly in external validation cohorts where some genes failed to reach significance thresholds, which may be related to insufficient sample size. Future studies should incorporate larger-scale multi-center cohorts for validation to improve the reliability and generalizability of results.

Second, there is a lack of functional validation experiments. This study is a retrospective bioinformatics investigation, and although it identified key genes and signaling pathways associated with PCOS, whether the expression changes of these genes are causes or consequences of the disease still requires functional experimental validation. Future studies should employ molecular biology approaches such as gene knockout/knockdown and overexpression in cell models (such as human granulosa cell line KGN, endometrial stromal cells) and animal models (such as DHEA-induced PCOS mouse models[38]) to validate the causal roles of key genes in PCOS pathogenesis. In particular, attention should be focused on the functions of CD44 in follicle development, inflammatory responses, and insulin resistance, as well as the effects of candidate drugs on improving PCOS phenotypes.

Third, transcriptome data cannot directly reflect protein levels and functional states. Changes in gene expression do not necessarily translate into corresponding changes at the protein level, especially for proteins regulated by post-translational modifications. For example, the activity of many signal transduction proteins is primarily regulated by post-translational modifications such as phosphorylation and ubiquitination, which cannot be captured by transcriptome data. Additionally, although molecular docking analysis can predict drug-target binding modes, it cannot reflect the true intracellular environment (such as pH, ion concentrations, competitive binding by other proteins). Therefore, integration with proteomics, phosphoproteomics, and metabolomics data for multi-omics analysis is needed, along with validation of drug-target interactions through in vitro binding experiments (such as surface plasmon resonance, isothermal titration calorimetry[39]).

Fourth, this study did not incorporate treatment intervention data. Whether the identified key genes and candidate drugs can serve as therapeutic targets, and whether interventions targeting these targets can improve clinical outcomes in PCOS patients, still requires further investigation. Future studies could explore therapeutic strategies targeting Hub genes such as CD44 and GJA5, evaluating the therapeutic effects of candidate drugs (such as Flufenamic acid, Cytosporone B) in preclinical models. Additionally, prospective cohort studies should be conducted, incorporating more detailed clinical information (such as alcohol consumption, BMI, HOMA-IR, AMH levels, oocyte retrieval numbers, pregnancy outcomes), to assess the associations between key gene expression patterns and clinical phenotypes and treatment responses, providing reference for stratified management and personalized intervention research in PCOS.

## 5. Conclusion

This study systematically characterized PCOS-related candidate genes and pathways by integrating differential expression analysis, weighted gene co-expression network analysis (WGCNA), PPI network analysis, and network pharmacology-based drug repurposing analysis. Based on three independent cohorts from the GEO database (GSE155489, GSE138518, GSE226146), an intersection strategy was employed to screen 498 core candidate genes from differentially expressed genes and significantly associated module genes. Functional enrichment analysis showed that, after multiple testing correction, candidate genes were significantly enriched in cell adhesion-related processes and TGF-beta signaling pathway entries, suggesting that PCOS granulosa cells may exhibit alterations related to intercellular communication and signal transduction.

Protein-protein interaction network analysis identified CD44 as the core Hub gene, which has the highest connectivity in the PPI network (degree=42) and centrality, and was significantly downregulated in differential expression analysis (logFC=-1.073, adj.P=0.00831). As the primary receptor for hyaluronic acid, CD44 may participate in follicle development, granulosa cell function regulation, insulin resistance-related signaling pathways, and endometrial receptivity regulation, serving as a candidate key molecule connecting the "ovary-metabolism-endometrium" integrated pathological process. This finding provides a new perspective for PCOS mechanistic research, suggesting that CD44 can serve as a potential target of interest for subsequent studies.

Drug repurposing analysis based on the CLUE platform (Connectivity Map/LINCS database) screened 106 candidate drugs, providing screenable clues for subsequent mechanistic research and candidate intervention directions. It should be emphasized that these results belong to computational inference based on expression reverse matching, and whether they have translational value still depends on validation across multiple aspects including tissue specificity, dose-response relationships, and safety.

Computational predictions from molecular docking and ADMET properties further provided screening criteria at the structural and physicochemical levels: for example, GJA5 with Flufenamic acid showed relatively stronger predicted binding affinity (-7.88 kcal/mol), and Cytosporone B had more ideal druglikeness indicators (0 Lipinski violations). It should be emphasized that these results are primarily used for prioritization and hypothesis generation, and still require further validation through in vitro/in vivo experiments and clinical evidence.

The main features of this study include: (1) employing an intersection strategy (DEGs ∩ WGCNA module genes) to narrow the candidate pool and combining network analysis to provide systems-level interpretive clues; (2) conducting supplementary verification of expression directions of key genes across multiple cohorts; (3) integrating drug repurposing results with molecular docking and ADMET computational predictions to provide a consistent analytical chain for prioritizing candidate targets and candidate molecules. Overall, this study can provide traceable candidate lists and analytical frameworks for PCOS-related mechanistic research and subsequent validation experiments.

## 8 CRediT authorship contribution statement

Xuan Zhang: Conceptualization, Methodology, Software, Investigation, Writing - Original Draft.

Jie Fang: Investigation, Data Curation, Validation.

Zhinan Liu: Formal analysis, Visualization, Resources.

Shuangsang Li: Methodology, Investigation.

Fanhui Jin: Validation, Data Curation.

Linlin Guo: Resources, Software.

Ruonan Qiang: Investigation, Validation.

Yinan Zhu: Investigation.

Tingting Hou: Data Curation.

Jingjing Li: Formal analysis.

Yanfeng Liu: Conceptualization, Supervision, Project administration, Funding acquisition, Writing - Review & Editing.

All authors have read and approved the final manuscript.

## 10 Availability of data and materials

The datasets analyzed during the current study are available in the Gene Expression Omnibus (GEO) repository (https://www.ncbi.nlm.nih.gov/geo/) under accession numbers GSE155489, GSE138518, and GSE226146.

## 11 Declarations

Competing interests

The authors declare no competing interests.

Disclosure

The authors have no competing interests to declare.

Ethics approval and consent to participate

This study was conducted using publicly available data, further ethical approval was not required for this secondary data analysis.

Clinical Trial Registration Number: Not applicable.

Consent for publication: Not applicable.

## Acknowledgments

We would like to express our sincere gratitude to the Gene Expression Omnibus (GEO) database for providing the high-quality transcriptomics datasets (GSE155489, GSE138518, and GSE226146) that made this study possible. We also thank the teams behind the CLUE platform (Connectivity Map/LINCS), the STRING database, the AlphaFold Protein Structure Database, and the developers of the R software packages (DESeq2, WGCNA, clusterProfiler) and AutoDock Vina, whose tools were instrumental in our bioinformatic and network pharmacology analysis. Finally, we thank all the researchers who shared their raw data and findings in the public domain, contributing to the advancement of PCOS research.

---

**Supplementary Figure S1A.**
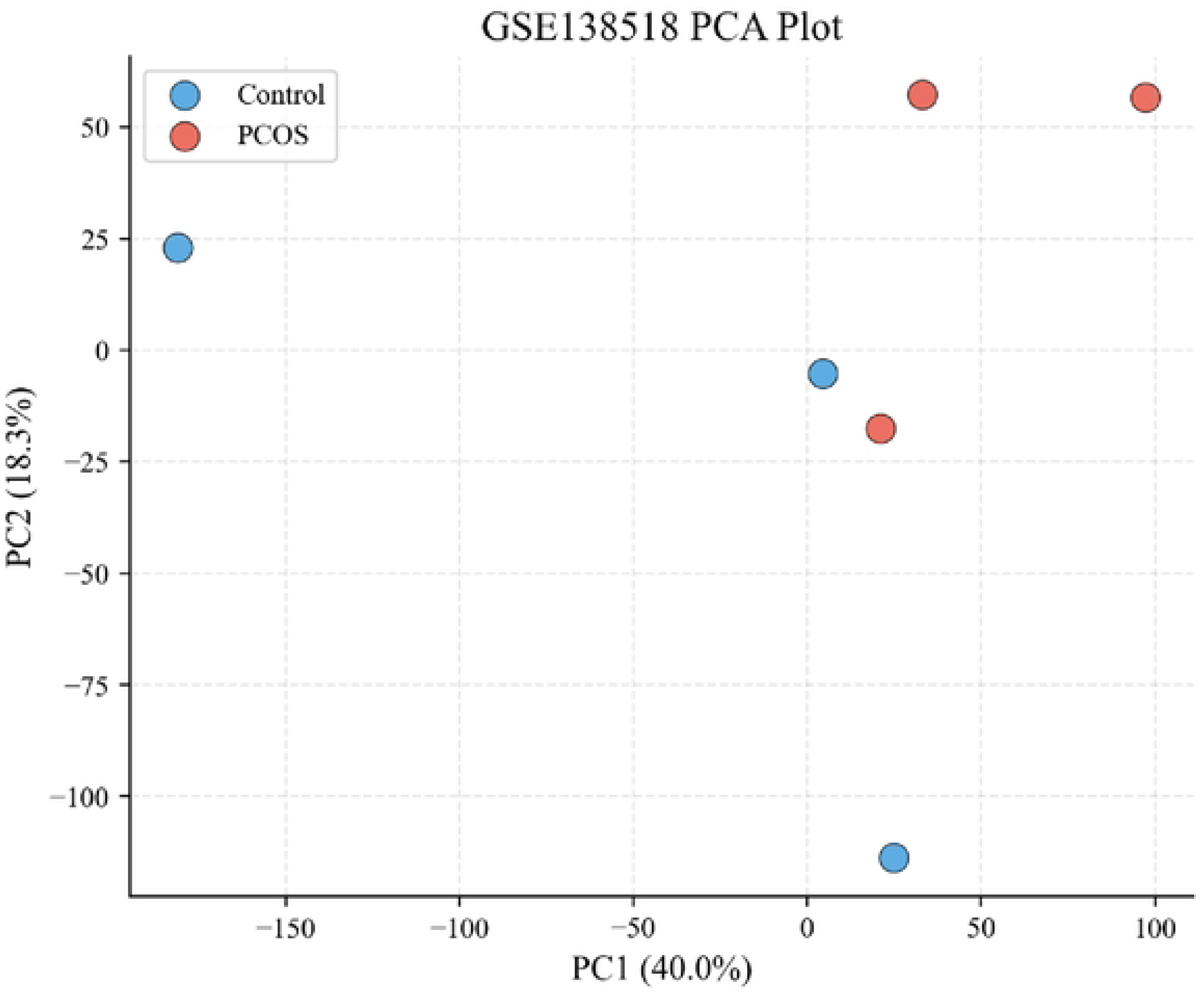
Principal component analysis of GSE138518 validation cohort (granulosa cells) Principal component analysis (PCA) showing overall transcriptomic differences between PCOS patients (n=3) and healthy controls (n=3) in the GSEl385 I8 dataset. This dataset is derived from ovarian granulosa cells, representing thesame tissue type as the discovery cohort GSEI55489. PCA was performed based on expression profiles of all genes, displaying sample distribution across the first two principal components (PC1 and PC2). The plot shows a degree of clustering by disease status, although the separation between groups is less pronounced than in the discovery cohort due to the small sample size (n=3 per group).

**Supplementary Figure S1B.**
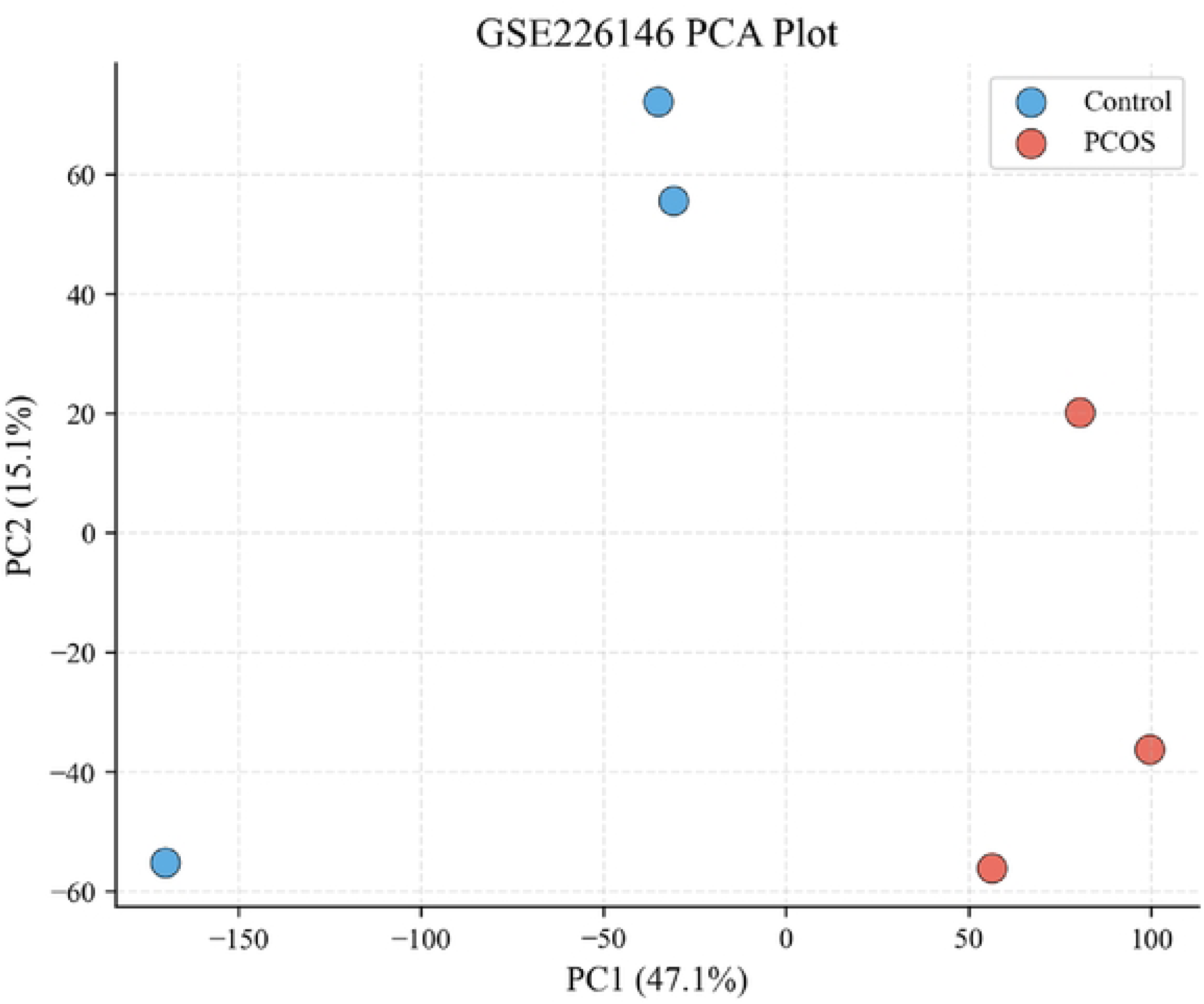
Principal component analysis of GSE226l 46 validation cohort (endometrium) Principal component analysis (PCA) showing overall transcriptomic differences between PCOS patients (n=3) and healthy controls (n=3) in the GSE226146 dataset. This dataset is derived from endometrial tissue, representing a different reproductive tissue type compared to the discovery cohort. PCA was performed based onexpression profiles of all genes, displaying sample distribution across the first two principal components (PCI and PC2). The plot shows clear separation between PCOS and control samples, suggesting that PCOS-relatcd transcriptional changes may extend beyond the ovarian microenvironment to other reproductive tissues. Despite the limited sample size (n=3 per group), this cohort demonstrated higher directional consistency of hub genes (91%) compared to the same-tissue validation cohort GSE!385 l8, suggesting that some PCOS-related transcriptional alterations may have systemic characteristics across tissues.

